# Topological design principle for the robustness of necroptosis biphasic, emergent, and coexistent (BEC) dynamics

**DOI:** 10.1101/2023.01.23.525173

**Authors:** Fei Xu, Xiang Li, Rui Wu, Hong Qi, Jun Jin, Zhilong Liu, Yuning Wu, Hai Lin, Chuansheng Shen, Jianwei Shuai

## Abstract

Biphasic dynamics, the variable-dependent ability to enhance or restrain biological function, is prevalent in natural systems. Accompanied by biphasic dynamics, necroptosis signaling dominated by RIP1 also appears emergent and coexistent dynamics. Here, we identify the RIP1-RIP3-C8 incoherent feedforward loop embedded with positive feedback of RIP3 to RIP1 is the core topology, and the scale-free feature of RIP3 peak value dictates necroptosis BEC dynamics. Entropy production is introduced to quantify the uncertainty of coexistent dynamics. RIP3 auto-phosphorylation is further determined as a complementary process for robustly attaining necroptosis BEC dynamics. Through screening all possible two- and three-node circuit topologies, a complete atlas of three-node circuit BEC dynamics is generated and only three minimal circuits emerge as robust solutions, proving incoherent feedforward loop is the core topology. Overall, through highlighting a finite set of circuits, this study yields guiding principles for mapping, modulating, and designing circuits for BEC dynamics in biological systems.

## Introduction

Biphasic behavior has been observed in a broad range of biological processes to drive essential physiological and developmental functions, such as cell differentiation^1^, proliferation^2^, and death^3^. Biphasic behavior can be broadly categorized into time-dependent and dose-dependent. Time-dependent biphasic behavior means the output response (*e.g.*, gene expression, protein activation, and ionic concentration, etc.) is increased with time at initial stage but becomes decreased over time. Conversely, the initial decrease and later increase could also be regarded as time-dependent biphasic behavior. Pulse and adaption are the typical time-dependent biphasic behaviors^4–6^, such as the transient ERK activation induced by growth factors^6^ and the PhoQ activation induced by ADP affinity in bacterial two-component system^7^. Dose-dependent biphasic behavior refers to that the output increases (or decreases) first and then decreases (or increases) with the increase of the input, such as the biphasic dose dependence on Norepinephrine in cyclic AMP signaling^8^, and the blue-light-dependent phosphorylation of Arabidopsis cryptochrome 2 in HEK293 cells^9^.

Biphasic behavior in biological systems that crosses the tipping point frequently accompanies by the emergence of new patterns. Emergent dynamic is the biological function outcomes of collective interaction among various components, such as the patterns of chimera states and synchronization triggered by cell-to-cell interactions^10, 11^. Most recently, we found RIP1 biphasically regulates RIP3 phosphorylation with necroptosis emergence in TNF-induced cell death signaling^12^. However, how the signaling topology is intrinsically related to RIP3 biphasic dynamics with emergence and how the biphasic dynamics are regulated, have not been elucidated. Exploring the link between biological functions and the design principles of biological networks is a fundamental challenge to understand how living organisms can perform various functions efficiently and accurately. The theories of network science have been proven to be powerful tools^13–15^. Despite the apparent complexity and diversity of cell signaling, only a limited number of topologies might be capable of robustly executing particular function. Ma et al. successfully dissected the principle for the design of network topologies that robustly achieve adaptation^4^. Li et al. validated that the robustness of biological oscillators is enhanced by the incoherent inputs^16^. The design principle for robust oscillatory behaviors with respect to noise also have been demonstrated recently^17^. Thus, revealing the properties of how biphasic and emergent dynamics are controlled and tailored in natural systems are urgently needed as well for understanding and optimizing biological regulatory strategies.

Starting with searching the essential structure to achieve RIP3 biphasic dynamics with necroptosis emergence, a TNF-induced death circuit model that well reproduces the experimental observations is proposed. The RIP1-RIP3-C8 incoherent feedforward loop is determined to be the core topology for biphasic dynamics with emergence induction, while the positive feedback of RIP1 activated by RIP3 dominates the coexistence of necroptosis and apoptosis. A scale-free feature of RIP3 peak value and the Bell-shaped regulation of RIP3 biphasic dynamics are further identified and analyzed. Previous study suggested that scale-free networks are empirically rare^18^, and the scale-free feature of RIP3 phosphorylation might be highly related to our recently determined composition of RIP1-RIP3 signaling hub^19^. To quantify the uncertainty of the coexistent dynamics, entropy production of the system is measured through introducing potential landscape and Shannon entropy theories for the first time. Instead of exploring the mechanisms in a specific system, random parameter analysis of the TNF circuit model is also performed, confirming the biphasic dynamics with emergence is the intrinsic properties of the death signaling topology. Besides the positive feedback of RIP1 activated by RIP3, the positive feedback of RIP3 self-activation embedded within the RIP1-RIP3-C8 incoherent feedforward loop is another fundamental structure for achieving the biphasic, emergent, and coexistent (BEC) dynamics. Finally, an exhausting search of all possible two- and three-node network topologies is performed to identify those capable of biphasic dynamics with emergence, and three categories of minimal circuits are obtained. Based on the minimal circuit, an optimal circuit structure for robustly achieving BEC dynamics is further proposed, which is highly consistent with the experimentally observed RIP1-RIP3-C8 circuit. Overall, all the evidence we obtained in this study indicates that the incoherent feedforward loop embedded with positive feedback is a generalizable design principle for the induction of BEC dynamics in diverse biological systems.

## Results

### Necroptosis BEC dynamics within TNF-induced death circuit

TNF is a multi-functional cytokine that can induce apoptosis or necroptosis depending on cellular contexts^20–22^. The schematic diagram of TNF-induced apoptosis and necroptosis signaling pathway is shown in Figure 1A. To intuitively address the relation of the core module, the reactions, such as association/disassociation, and cascades reaction can be coarsely described to present a conceptual core signaling circuit as shown in Figure 1B. Upon stimulation, TNF combines with TNFR1 to recruit TRADD and RIP1 to form complex-I and then activates them (Figure 1A)^23^, which can be simplified as TNF activating TRADD and RIP1 in Figure 1B. The competition between TRADD and RIP1 for binding TNFR1 is described as mutual inhibition^24^. Activation of C8 by sufficient TRADD in complex-II (labeled as C8-IIa) could result in apoptosis. When apoptosis occurs, the cell volume becomes small and the nucleus shrinks^25^. Besides, C8 could also be activated by RIP1 in necrosome (labeled as C8-IIb)^26^. In necrosome, C8 inhibits the phosphorylation of RIP1 and RIP3 through cleaving RIP1-RIP3 complex^27^. Phosphorylation of RIP3, the marker of necroptosis^21^, also blocks C8 activation by recruiting RSK^28^. RIP1 and RIP3 activate each other through their RHIM-domain, forming a positive feedback loop for the recruitment of MLKL and necroptosis induction^29, 30^. As a result, the TNF-induced death dynamics are mainly determined by five components, *i.e.*, activated TRADD (acTRADD), phosphorylated RIP1 (pRIP1), phosphorylated RIP3 (pRIP3), C8 activated by TRADD (C8-IIa), and C8 activated by RIP1 (C8-IIb) (Figure 1B).

As the western blotting data shown in Figure 1C, RIP1 knockdown (RIP1 shRNA) accelerates RIP3/MLKL phosphorylation, while RIP1 deletion (RIP1 KO) completely blocks RIP3/MLKL phosphorylation, presenting a biphasic dynamics of necroptosis regulated by RIP1. Decrease of RIP1 also promotes C8 activation, suggesting the suppression role of RIP1 in apoptosis (Figure 1C). To fully understand the essential topology for achieving the biphasic dynamics in the death signaling, a self-evolving ODEs model is constructed based on the circuit shown in Figure 1B (Supplementary Text). The circuit model well reproduces the experimental observations of C8 and RIP3 activation under different RIP1 levels (Figures 1D and 1E). pRIP3 presents an abrupt and large increase at low level of RIP1, triggering the emergent dynamics of necroptosis (Figure 1E). pRIP3 is then gradually reduced with further increase of RIP1. While C8 activation is linearly reduced with RIP1 increase. Experiments show that the deletion of RIP1-induced C8 activation is completely blocked in TRADD deletion cells (Figure 1F), proving that RIP1 suppresses apoptosis through restraining TRADD-dependent C8 activation. Consistently, our circuit model can also quantitatively reproduce the experimental observations and further provides a comprehensive analysis result, showing the decrease of TRADD results in a progressive reduction of C8 activation at varying RIP1 levels (Figure 1G).

As RIP3 and C8 can be simultaneously activated (Figure 1C), coexistence death mode of apoptosis and necroptosis in cells could be triggered by proper RIP1 level. Experimental analysis of cell morphology suggests that only necroptosis occurs in wild-type (WT) cells (100% RIP1), while apoptosis can solely be observed in RIP1 deletion cells (0% RIP1) (Figure 1H, upper panel). As expected, both necroptosis and apoptosis can be observed in RIP1-impaired cells. The RIP1-induced cell death mode can be well described by potential landscape theory, which provides a more physical description of the stochastic dynamic and global stability of the biological system^31–33^. Consistent with experimental observations, the middle panel from left to right in Figure 1H are the landscape topography of RIP1 deletion (single apoptosis state), RIP1 impairment (coexistent state of apoptosis and necroptosis), and WT (single necroptosis state) systems that are mapped in the C8-RIP3 phase space. The kinetic pathway (KP) of the system evolving from random initial state (RIS) in the bottom panel of Figure 1H visually presents how the decision of death mode is made under different RIP1 levels. With low or high RIP1 level, cells eventually have a unique death mode of apoptosis or necroptosis. While cells have two mode choices with proper RIP1 level, inducing the coexistent dynamics of apoptosis and necroptosis. Thus, above comparisons confirm that our circuit model has the potential for giving mechanistic insights into the pRIP3/necroptosis BEC dynamics within the TNF-induced death signaling.

**Figure 1.**
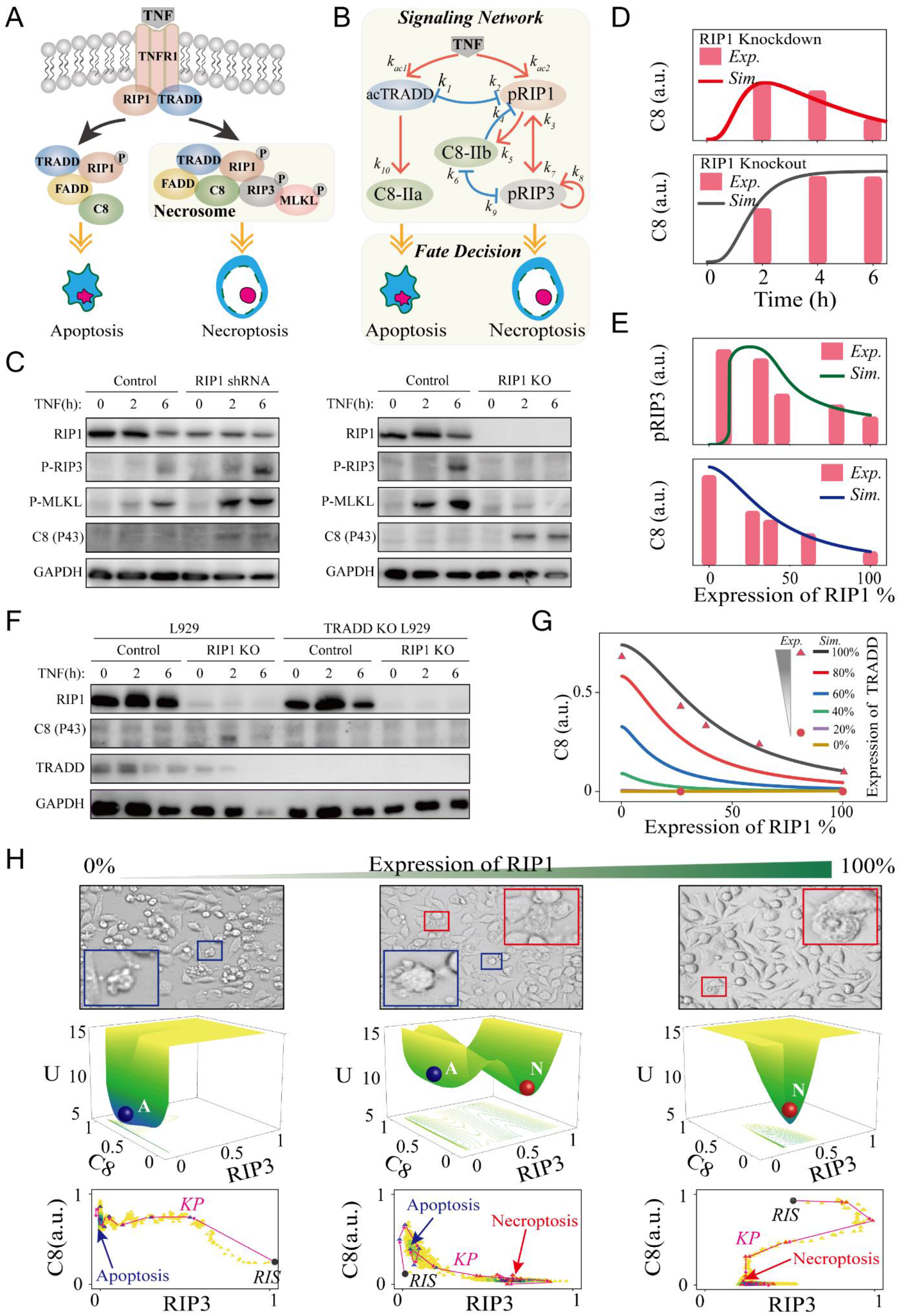
Data-driven modeling of the TNF-induced cell death circuit. (A) Schematic diagram of TNF-induced apoptosis and necroptosis signaling pathway. (B) The coarse-grained signaling network model. (C) Western blot analysis of the effects of RIP1 knockdown (shRNA) or knockout (KO) on indicated proteins activation. (D) Comparison between experimental data (histograms) and simulation results (lines) of the time-course responses of C8 in RIP1 knockdown (upper panel) and RIP1 knockout (down panel) cells. (E) Comparison between experimental data (histograms) and simulation results (lines) of RIP1-dependent pRIP3 response (upper panel) and C8 activation response (down panel). (F) Western blot analysis of TRADD knockout on indicated protein activation for wildtype and RIP1 knockout cells. (G) Comparison between experimental data (dots) and simulation results (lines) of the effect of TRADD level on RIP1-dependent C8 activation. (H) Cell morphologies under different expression levels of RIP1. The red and blue boxes indicate the represented apoptotic and necroptotic cells, respectively. Potential landscape reveals switch in cell death modes under different levels of RIP1 and the corresponding kinetic pathways (KP) of system evolution from random initial states (RIS).

### RIP1-RIP3-C8 incoherent feedforward loop determines necroptosis BEC dynamics

To dissect the essential topology for the BEC dynamics of pRIP3 induced by RIP1, roles of TRADD and C8 are first explored. Biphasic and emergent (BE) dynamics of pRIP3 are not affected by TRADD deletion (Figure 2A, blue line), but disappear in the absence of C8 (Figure 2A, green line), implying that C8 is the essential node for BE dynamics. Deletion of C8 barely affects the emergence of pRIP3, and pRIP3 level keeps constant with further increase of RIP1. Thus, the dynamics of pRIP3 consists of two processes: when RIP1 increases from 0 to a low critical level (∼10% RIP1), pRIP3 increases abruptly, inducing the emergent dynamics of necroptosis. While with the increase of RIP1, inhibition of C8 on pRIP3 takes effect, causing pRIP3 gently decreases.

Then, the nine interaction terms among RIP1, RIP3, and C8 in the irreducible circuit model are respectively removed to determine the essential terms (Figure 2B). BE dynamics disappear when the terms including *k*_5_ (C8 activated by RIP1), *k*_7_ (RIP3 activated by RIP1), and *k*_9_ (RIP3 inhibited by C8) are respectively set to 0, while BE dynamics are still observed when the other six terms are fixed to 0. We introduced the coefficient H, which is defined as *H* = (*pRIP3_Peak_*-*pRIP3_RIP1_100%_*) / *pRIP3_tot_* to quantify the scale of biphasic dynamics. *pRIP3_Peak_* is the maximum level of pRIP3, and *pRIP3_RIP1_100%_* is the level of pRIP3 when the expression level of RIP1 is 100% (wild-type). Therefore, analysis in Figure 2B indicates that the essential topology for necroptosis BE dynamics of the TNF-induced death circuit is constituted by the incoherent feedforward loop structure (*k*_5_, *k*_7_, and *k*_9_) that are embedded in the three nodes (RIP1, RIP3, and C8).

Besides the identified RIP1-RIP3-C8 topological structure for BE dynamics, the interaction term required for achieving the coexistence mode of necroptosis and apoptosis is further explored based on potential landscape analysis. We respectively removed the terms besides the identified essential topology for BE dynamics to determine whether apoptosis and necroptosis states could coexist with RIP1 variation. As shown in Figure 2C, only when the term of RIP1 activated by pRIP3 (*k*_3_) is removed (Figure 2Ciii), the system presents solely one potential well with high C8 and low pRIP3 level. While the system still exhibits two coexisting wells with the blockage of other terms. Taken together, as the diagram shown in Figure 2D, RIP1-RIP3-C8 incoherent feedforward loop embedded with the positive feedback of RIP3 to RIP1 is the core topological structure for achieving RIP1-induced necroptosis BEC dynamics.

**Figure 2.**
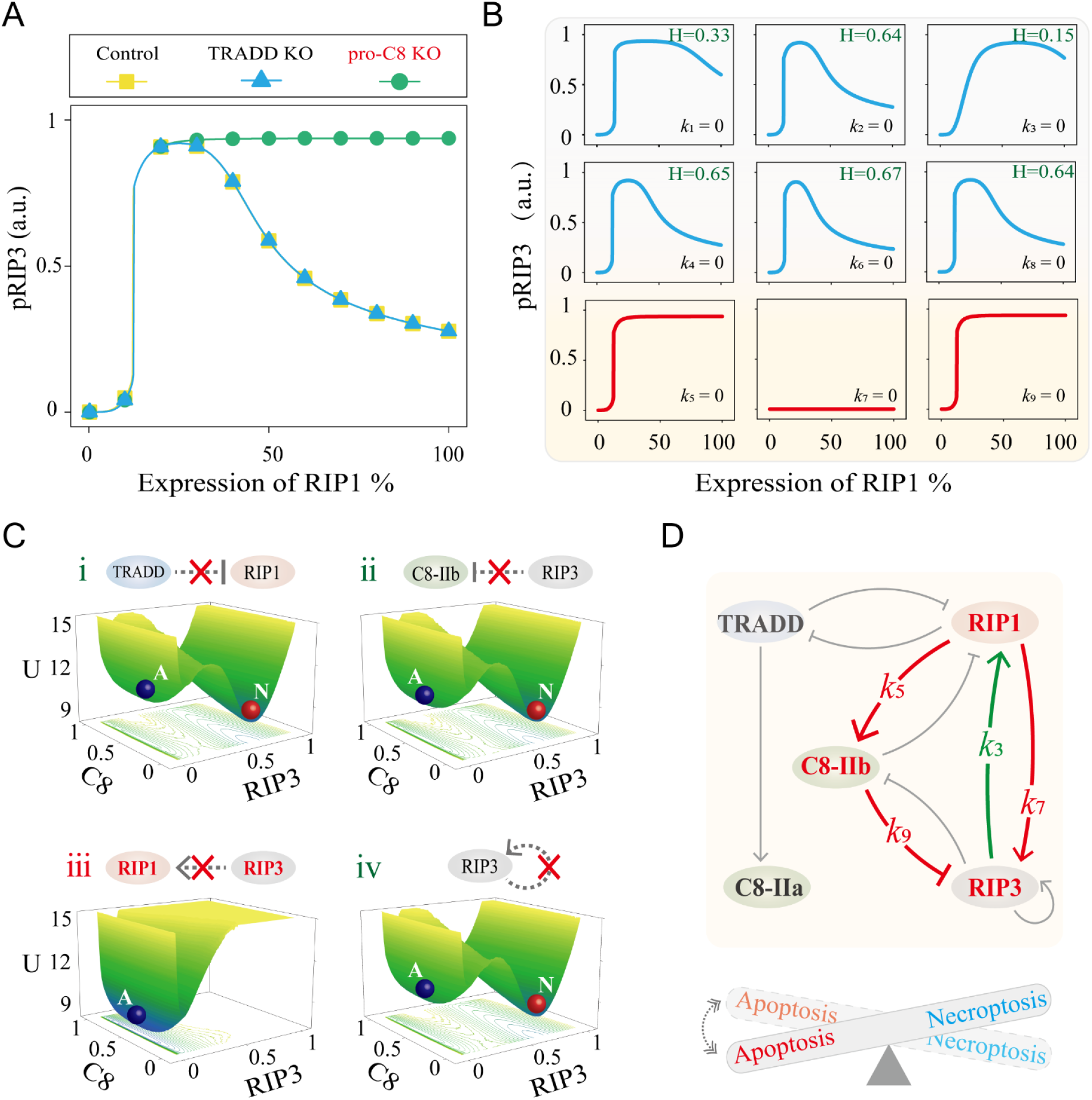
Identification of the core structure for pRIP3/necroptosis BEC dynamics. (A) Comparison of pRIP3 BE dynamics under control, TRADD KO, and C8 KO conditions. (B) The dynamics of pRIP3 when any one of the nine interaction terms in circuit model is removed, respectively. (C) Potential landscapes of the system when the four interaction terms are severally removed. (D) Summary of the constituents and terms that can achieve BE dynamics (red lines) with coexistent death mode (green line).

### Bell-shaped regulation of necroptosis biphasic dynamics by RIP3 and the feedforward terms

Having identified the essential topological structure, we next explored the control mechanism of how necroptosis biphasic dynamics is generated and regulated by the protein nodes (RIP3, TRADD, and C8) and interaction terms. Both the absolute and relative levels of *pRIP3_Peak_* and *pRIP3_RIP1_100%_* to RIP3 expression level variation are shown in Figure 3A. The absolute level of *pRIP3_Peak_* is linearly positively correlated with RIP3 (upper left panel of Figure 3A), whereas the relative level of *pRIP3_Peak_* (*pRIP3_Peak_/RIP3_tot_*) remains constant, presenting a scale-free feature (upper right panel of Figure 3A). For *pRIP3_RIP1_100%_*, the absolute level is also positively related to RIP3, but the relative level exhibits a biphasic behavior (down panel of Figure 3A). The scale-free feature of *pRIP3_Peak_* and biphasic behavior of *pRIP3_RIP1_100%_* result in the biphasic dynamics of RIP3 presents inverted Bell-shaped responses to RIP3 expression level, as the quantified scale of biphasic dynamics H shown in Figure 3B. Four-Parameter Logistic Function is considered for a piecewise fit of H. When RIP3 is lower than ∼10%, H decreases monotonically from the maximum value of 0.44 to 0.18, and the decline rate is the largest at ∼5%. Conversely, H is positively correlated with RIP3 when RIP3 is higher than ∼10%. The fitted function suggests that the maximum value of H cannot exceed 0.72 with a maximum increase rate at ∼40%.

To systematically reveal the Bell-shaped regulation mechanism of RIP3 on the scale of H, the variation of relative pRIP3 (*pRIP3*/*RIP3_tot_*) is investigated in RIP3-RIP1 phase plane (Figure 3C). The plane can be divided into 5 regions with specific regulatory mechanisms. Region 1 indicates that the inhibition of TRADD on RIP1 is dominant and RIP3 remains inactive with low expression level of RIP1 (Figure 3Di). When RIP1 increases to a critical level, RIP1 activated RIP3 and the self-activation of RIP3 induce the emergence of pRIP3 (Figure 3Dii), corresponding to region 2. Regions 3, 4, and 5 show that high level of RIP1 negatively regulates pRIP3. In region 3 with the low expression level of RIP3, pRIP3 is greatly restrained by C8 (Figure 3Diii), exhibiting a large value of H. As the expression level of RIP3 increases in region 4, inhibition of pRIP3 on C8 gradually becomes dominant (Figure 3Div). pRIP3 level increases and H decreases. Further increase of RIP3 in region 5 enhances the positive feedback of pRIP3 on RIP1, which indirectly promotes the inhibition of C8 on pRIP3 (Figure 3Dv), resulting in a resurgence of the large scale of biphasic dynamics (large value of H). We confirmed the inferences though respectively reducing the strength of the corresponding interaction terms, which characterize the mutual inhibition between C8 and pRIP3, and the positive feedback of pRIP3 on RIP1 (Figure S1A). Specifically, when the strength of C8 inhibition on RIP3 (*k*_6_) decreases, region 3 disappears and the relative level of pRIP3 in region 5 increases. While weakening the inhibition strength of pRIP3 on C8 (*k*_9_) results in the disappearance of region 4 and the expansion of region 3. Attenuating the positive feedback strength of pRIP3 on RIP1 (*k*_3_) increases the relative level of pRIP3 in region 5. The scale of biphasic dynamics H is barely influenced by TRADD (Figure S1B), but is gradually enhanced with the increase of C8 (Figure S1C). For C8, the relative *pRIP3_Peak_* also presents a scale-free feature, while the relative *pRIP3_RIP1_100%_* is linearly decreased with the increase of C8.

We next investigated the role of interaction terms in mediating the scale of biphasic dynamics H. Among the nine terms, only *k*_5_ (C8 activated by RIP1) and *k*_7_ (RIP3 activated by RIP1) that involves in the RIP1-RIP3-C8 incoherent feedforward loop, can both mediate the levels of *pRIP3_Peak_* and *pRIP3_RIP1_100%_*, achieving Bell-shaped regulation on H (Figure 3E and S2). With the increase of *k*_5_, *pRIP3_Peak_* decreases first slow and then fast, while *pRIP3_RIP1_100%_* decreases first fast and then slow, presenting distinct responses (Figure 3E). In contrast, increase of *k*_7_ makes *pRIP3_Peak_* to increase first fast and then slow, but *pRIP3_RIP1_100%_* to increase first slow and then fast. The scale H is the largest when the change rate of *pRIP3_Peak_* and *pRIP3_RIP1_100%_* are equal with the corresponding strengths *k*_5_=10^-0.24^ and *k*_7_=10^0.28^. *pRIP3_Peak_* can hardly be regulated by the rest seven terms, while *pRIP3_RIP1_100%_* is positively regulated by *k*_6_ and negatively regulated by *k*_1_, *k*_3_, and *k*_9_ (Figure S2A). Further two-parameters phase plane analysis indicates that the maximum value of H is 0.68 when *k*_5_=0.575 and *k*_7_ =1.82 (Figure 3F). We severally profiled *pRIP3_Peak_* and *pRIP3_RIP1_100%_* in the *k*_5_-*k*_7_ phase plane (Figure S3A), and three different regions and two processes are identified in the plane to reveal the mechanism of Bell-shaped regulation (Figure S3B). Although the term of *k*_9_ (inhibition of C8 on pRIP3) that involves in the RIP1-RIP3-C8 incoherent feedforward loop can not drive Bell-shaped regulation on H, *k*_9_ significantly amplifies the Bell-shaped regulation of *k*_5_ (C8 activated by RIP1) or *k*_7_ (RIP3 activated by RIP1) on H (Figure S3C).

**Figure 3.**
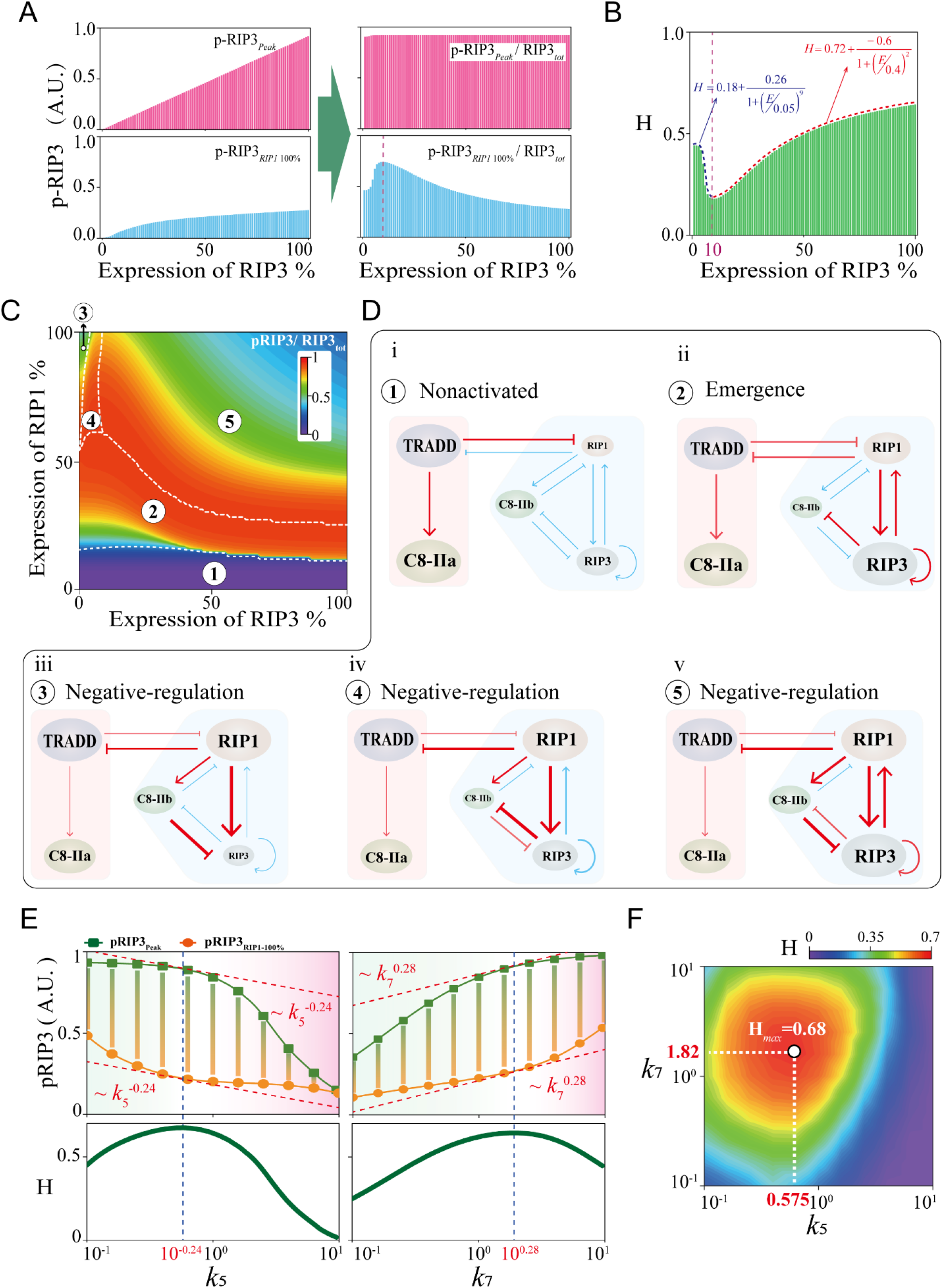
Scale-free emergence of pRIP3 and Bell-shaped regulation of necroptosis biphasic dynamics. (A) The variation of absolute and relative levels of *pRIP3_Peak_* and *pRIP3_RIP1_100%_* with RIP3 expression level increases. (B) The quantified scale H of pRIP3 biphasic dynamics. (C) The relative level of pRIP3 in the RIP3-RIP1 phase plane, giving the plane divided into 5 regions. (D) Topological analysis of the regulatory mechanisms of the 5 regions in (C). (E) Analysis of the *k*_5_ and *k*_7_ Bell-shaped regulation on pRIP3 biphasic dynamics. (F) Phase diagram of H in *k*_5_-*k*_7_ parameter spaces.

### Shannon entropy quantifies uncertainty of the coexistence death modes

Previous studies attempted to infer the information for thermodynamic quantities by observing dwell time distribution of transitions among multiple steady states^34–37^. The cell death signaling presents coexistent dynamics with proper RIP1 level, as the two basins (apoptosis and necroptosis) shown in the death landscape topography (Figure 1G). To measure the uncertainty of cell fate decision, we introduced Shannon entropy to estimate entropy production, which is defined as *S* = – Σ*p_i_log*_2_(*p_i_*). *p_i_* is the probability of the *i*-th death mode, and *i* represents the state of apoptosis or necroptosis. S characterizes the degree of disorder of the cell death system. The larger the value of S is, the more disorder the system is. Figure 4A illustrates the analysis procedure for Shannon entropy by calculating dwell time distribution. Stochastic cell death system is obtained by adding Gaussian white noise to the deterministic system with random initial states. Temporal dynamics of the system are precisely recorded at the free degrees of pRIP3 to obtain the dwell time distribution of the system in different states. With the increase of RIP1, the dwell time in apoptosis state is gradually decreased, but is increased in necroptosis state (Figure 4B, upper panel), and the corresponding dwell time distributions are shown as well (Figure 4B, down panel).

Shannon entropy of *k*_5_ (term of C8 activated by RIP1 that exhibits Bell-shaped regulation on necroptosis biphasic dynamics) in regulating RIP1-dependent coexistent dynamics is measured (Figure 4C). The region surrounded by the white dotted line is the coexistence transition region. The color code in the transition region indicates the degree of transition between apoptosis and necroptosis of the system. The system presents a highly disordered state when RIP1 is near the level to induce pRIP3/necroptosis emergent dynamics. Shannon entropy (S) gradually decreases and necroptosis becomes dominated with further increase of RIP1. Thus, the cell fate switches from ordered apoptosis to highly disordered and finally to ordered necroptosis with the increase of RIP1. A more intuitive potential landscape topography transition is shown in Figure 4D when *k*_5_ is fixed at the standard value of 0.3. With the increase of RIP1, the depth of the apoptosis basin is gradually decreased, while the necroptosis basin is increased, suggesting that RIP1 biases the cell fate towards necroptosis. The transitions of Shannon entropy and potential landscape topography with different *k*_5_ are also investigated with RIP1 is fixed at 11% (Figure 4E). In contrast to RIP1, increase of *k*_5_ results in the uncertainty of cell fate switches from highly disordered to ordered apoptosis, giving the depth of apoptosis basin increased and necroptosis basin decreased. The entropy result also reveals that the increase of *k*_5_ causes a high level demand for RIP1 to trigger pRIP3 emergent dynamics (white arrow in Figure 4C). The larger the value of *k*_5_ is, the higher RIP1 level is required for inducing necroptosis emergent dynamics.

Therefore, acting as the driving force, RIP1 makes the system dynamics like a “seesaw” (Figure 4F). The system exclusively executes apoptosis at low RIP1 level. Increase of RIP1 significantly elevates pRIP3 level and entropy production. The death system is highly disordered and will selectively undergo apoptosis or necroptosis, depending on the initial conditions. High level of RIP1 reduces entropy production and drives the system to ordered necroptosis. RIP1-dependent coexistent dynamics regulated by the other three reactions within the identified essential topological structure (Figure 2D) are shown in Figure S4, indicating that the terms of *k*_3_ (RIP1 activated by pRIP3) and *k*_7_ (RIP3 activated by RIP1) can efficiently switch death modes, while the system remains highly disordered with the variation of *k*_9_ (inhibition of C8 on pRIP3).

**Figure 4.**
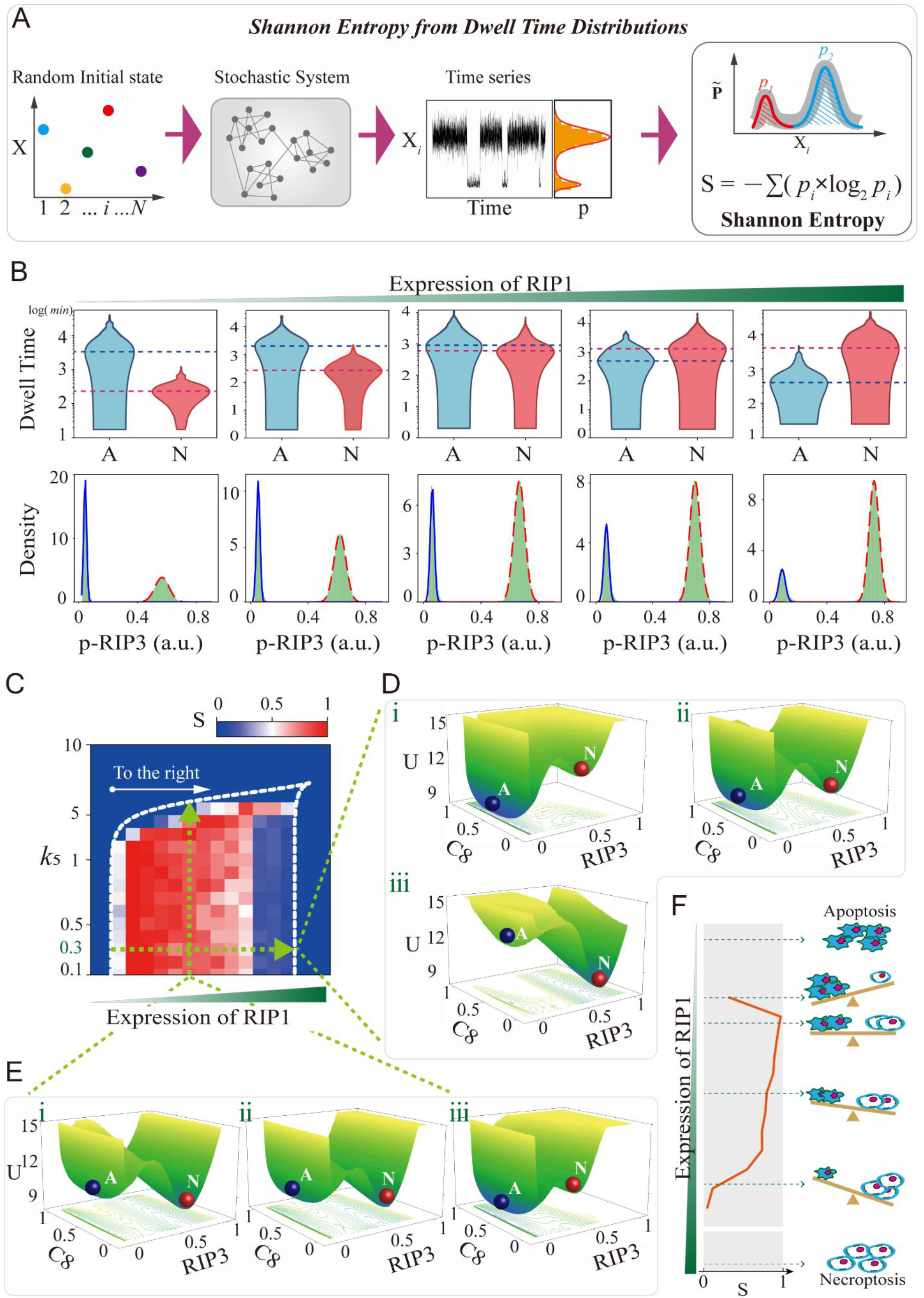
Shannon entropy quantifies the uncertainty of cell fate decisions. (A) Illustration of the calculation procedure of Shannon entropy with dwell time distribution. (B) Statistics and distribution of dwell time under five representative RIP1 expression levels at the free degrees of pRIP3. (C) The quantified Shannon entropy of k_5_ in regulating RIP1-dependent coexistent dynamics. (D) and (E) The potential landscape topography of coexistent death modes with k5=0.3, and RIP1 level at 10%, 11%, and 12.5% respectively (D), and the level of RIP1 at 11% with k5=0.1, 0.5, and 0.7 respectively (E). (F) Shannon entropy characterizes the uncertainty of cell fate, and a diagram of “seesaw” that reflects the death modes decision under different RIP1 levels.

### Random circuit analysis identifies two core topologies for necroptosis BEC dynamics induction

To avoid the fortuity of deterministic model with specific parameters, random circuit analysis is further performed to identify the core topology for necroptosis BEC dynamics. Latin hypercube sampling is used to obtain uniform random parameter regimes within reasonable biological interval^4, 12, 38^, and all parameter regimes that can achieve BE, and BEC dynamics are screened (Figure 5A). We assumed that the biphasic dynamics (BD) of pRIP3 satisfies *pRIP3_Peak_* is higher than *pRIP3_RIP1_100%_* by more than 1%, *i.e.*, BD is defined when BD=(*pRIP3_Peak_* – *pRIP3_RIP1_100%_*)/*pRIP3_Peak_* >=1%. Since pRIP3 presents an abrupt and large increase at low level of RIP1 (Figure 2A), emergent dynamics (ED) is also considered through satisfying pRIP3 is increased by more than 50% when RIP1 continuously increases by 10% (ED=Δ*pRIP3*/Δ*RIP1*>=5).

The probabilities under five representative RIP3 expression levels are firstly calculated, and 50,000 groups of random samples are taken for each RIP3 level. The probabilities of achieving BE dynamics (blue histograms) and BEC dynamics (green histograms) are presented in Figure 5B. As the result indicated, the vast majority (∼86.6%) of samples that achieve BE also have coexistent dynamics, revealing that the necroptosis BE dynamics is highly related to coexistence of death mode. The probabilities of BE and BEC dynamics decrease monotonically with RIP3 decreases. The probability of only considering biphasic dynamics is decreased by 17% when RIP3 decreases to 20% (Figure S5A). Whereas, the probability of BEC dynamics is decreased by 41.6% (Figure 5B). Thus, compared to biphasic, RIP3 seems to be more critical for the achievement of emergent and coexistent dynamics.

To determine the regulation of RIP3 on the scale of biphasic dynamics H, 570 samples that achieve BEC dynamics for 100% RIP3 are selected. The regulation of H by RIP3 has three types and their corresponding proportions are calculated: negative regulation (H is decreased with RIP3 increases) with 16.49%, positive regulation (H is increased with RIP3 increases) with 54.74%, and Bell-shaped regulation (nonlinear change in H with RIP3 increase) with 28.77% (Figure 5C). Three specific systems are further selected to intuitively show how RIP3 negatively, positively, or nonlinearly regulates H (Figure 5D, upper and middle panels). Another specific system is also selected to present the death mode transitions from apoptosis to the coexistence of apoptosis and necroptosis, and finally to the exclusive necroptosis state with the increase of RIP1 (Figure 5D, down panel).

Similar regulation of C8 on H is shown in Figure S5B, where the proportion of systems that achieving Bell-shaped regulation of H by C8 is less than ∼10% compared to the regulation of RIP3 on H (Figure 5C). Thus, variation of RIP3 seems to be more easily than C8 to achieve Bell-shaped regulation on H, supporting the results in the deterministic system that H does not present Bell-shaped response to C8 variation (Figure S1C). The average and variation of *pRIP3_peak_* with the decrease of RIP3 or C8 in the 570 systems are calculated as well (Figure 5E). The average of relative *pRIP3_peak_* is concentrated at a high level and the variation is quite small, revealing that the scale-free feature of necroptosis emergence (Figure 3A) is an intrinsic topological property of the death circuit.

The essential module for achieving BE and BEC dynamics are discussed with random circuit analysis as well. The interaction terms are completely blocked one by one in the 50,000 random samples, and the statistical probabilities are shown in Figure 5F. Only when any one of the terms (*k*_5_, *k*_7_, or *k*_9_) is removed, the probability tends to be 0, suggesting that the RIP1-RIP3-C8 incoherent feedforward loop is the necessary module to generate BE and BEC dynamics (Figure 5F and Figure S5C). Positive feedback loop is proven to be necessary for coexistent dynamics^39–41^. Unlike the deterministic system (Figure 2C), blocking the positive feedback of RIP3 to RIP1 (*k*_3_) cannot arrest coexistent dynamics (Figure 5F). While the probability of BEC is significantly decreased compared to BE with the blockage of RIP3 self-activation (*k*_3_). The contributions of all the four positive feedback loops within the death circuit are studied (Figure S5D), revealing that only the positive feedback loop formed by *k*_3_ or k_8_ is efficient for achieving the high occurrence probability of coexistent dynamics.

The regulation strategies of H by all the circuit interaction terms are separately calculated (Figures 5F and S5E). Consistent with deterministic system analysis (Figure 3E), the Bell-shaped regulation of H by the two feedforward terms of *k*_5_ and *k*_7_ are also statistically confirmed (Figure 5F). Moreover, RIP3 self-activation term *k*_8_ can also present Bell-shaped regulation on H, which is not observed in the deterministic system (Figure S2B). Thus, besides the positive feedback of RIP3 to RIP1 (Figures 5Gi and 2D), RIP3 self-activation forms another positive feedback for efficiently achieving coexistent dynamics and the Bell-shaped regulation on necroptosis biphasic dynamics (Figure 5Gii). The RIP1-RIP3-C8 incoherent feedforward loop embedded with these two positive feedback loops forms the fundamental hypermotif for robustly achieving BEC dynamics in the death circuit (Figure 5G).

**Figure 5.**
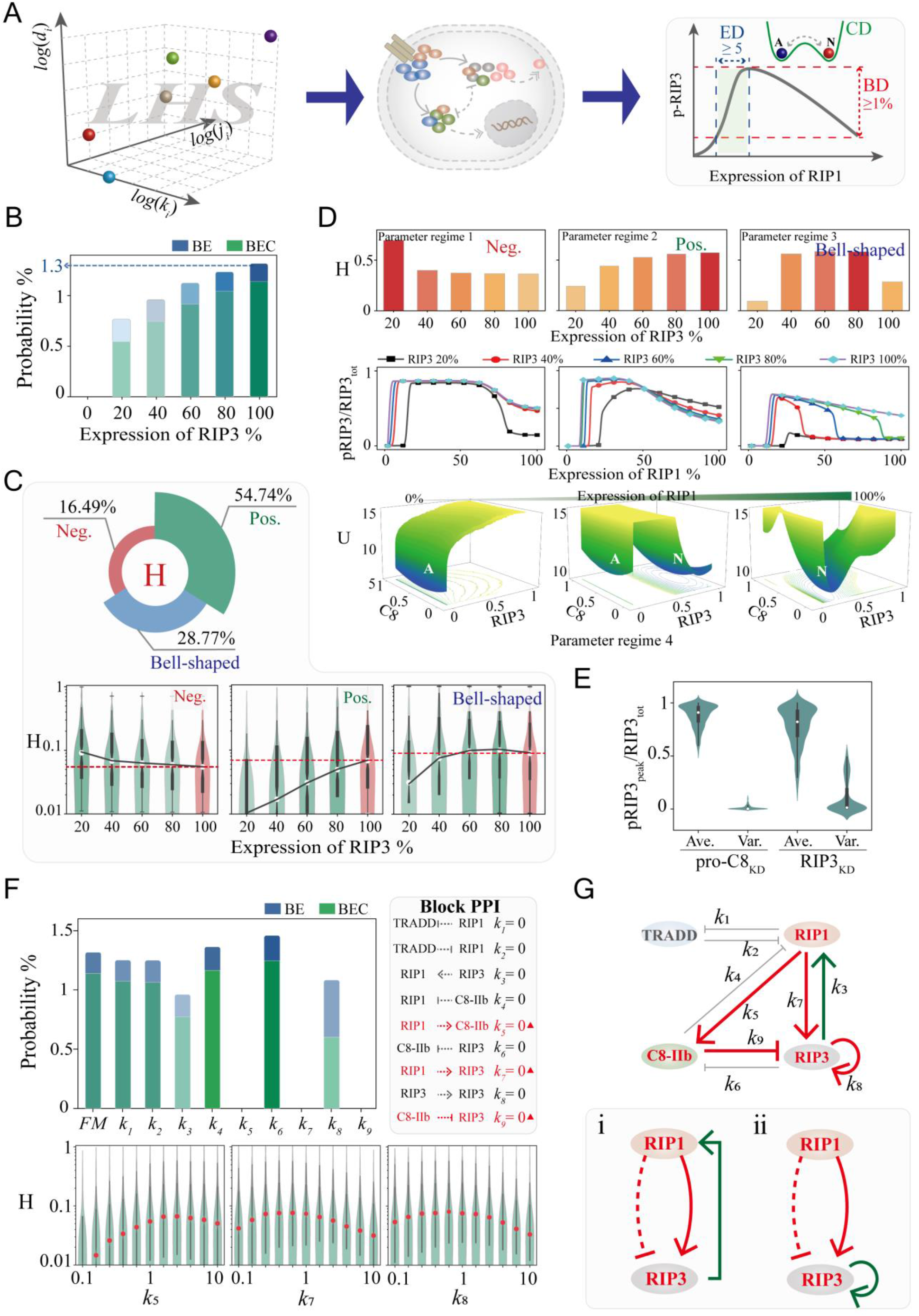
Identifying the core structure for necroptosis BEC dynamics with random circuit analysis. (A) Schematic of a computational workflow. Left panel: Latin hypercube sampling to obtain random parameter regimes. Right panel: Threshold settings for screened RIP1-dependent pRIP3 dynamics, including biphasic, emergent, and coexistent dynamics (BD, ED, and CD). (B) Random circuit analysis is used to search 50,000 systems under five representative RIP3 expression levels and count the probabilities that pRIP3 dynamic behaviors satisfy the threshold condition of BE and BEC dynamics. The blue and green histograms represent the probability of BE and BEC dynamics, respectively. (C) Statistics of the regulatory behavior of RIP3 on H in the screened systems with BEC dynamics. (D) Four representative systems of three different regulations of RIP3 on H (parameter regimes 1-3) and RIP1-dependent death modes switching behavior (parameter regimes 4). (E) The average and variance statistics of pRIP3peak relative level of all the screened samples with different expression levels of RIP3 and C8. (F) The probabilities of the system achieving BE (blue histograms) and BEC dynamics (green histograms) when any one term is removed. The terms of *k*_5_, k_7_, and k_9_ (marked by red triangles) are individually removed and the system cannot achieve biphasic dynamics. Down panels: Statistics of terms strength on H in the screened systems. (G) The components and reactions in the death circuit to achieve BEC dynamics. Red lines indicate the necessary interaction of the incoherent feedforward loop. Green lines indicate the positive feedback to RIP3.

### Topological exhaustivity defines three minimal circuits to achieve natural BE dynamics

To dissect the hidden design principles in biological systems, we tried to find the minimal circuit that performs biphasic dynamics with emergence. Topological exhaustive method has been widely used to explore the design of functional achievement in biological networks^4, 16, 42–44^. The workflow for circuit topology to function mapping of BE dynamics is shown in Figure S6A, which presents the circuit model described by coupling matrices, parameter regimes, and ordinary differential equations. We first examined whether the output signal node in the two-node circuit could achieve BE dynamics by the variation of the receiving node of input. All the 27 different structures of two-node circuit are respectively assigned 100,000 sets of random parameter regimes, but none of these circuits can achieve BE dynamics (Figure S6B).

We next searched BE dynamics in the three-node circuit, which includes an input node (node-A), an output node (node-C), and a regulatory node (node-B) (Figure 6A). There are three types of term (promotion, inhibition, and no interaction) between any two nodes. To map out the entire design space of three-node circuits capable of BE dynamics, all the 4,698 circuits are exhausted and analyzed with 50,000 sets of random parameter regimes assigned to each circuit. A total of 234.9 million dynamical systems (4,698×50,000 parameter regimes) are analyzed and finally 1,701 circuits that can achieve BE dynamics are screened out. To quantify the volume of the parameter space in which circuit supports BE dynamics, the 4,698 coupling matrices are firstly reduced to two-dimensional space using *t-SNE* method^45^, and then the probabilities are converted into topological potentials through -ln(*p*) analogy to the Boltzmann relation (Figure 6B). As a result, the three potential wells in the topological landscape correspond to three sub-clusters respectively. The three minimal circuits (M1, M2, and M3) in each sub-cluster are shown in Figure 6C, indicating that the circuits have the common feature of containing incoherent feedforward loop. Similar results are obtained with clustering analysis using the pair-wise distance between circuits, which also divides the 1,701 circuits into three sub-clusters and finally refers to the same three minimal circuits (Figure S7A).

Then, we severally segmented the obtained probabilities of the three sub-clusters into the two-dimensional space of biphasic and emergent dynamics (Figure 6D). Probability distributions of the three sub-clusters are quite different in the scale space. The sub-cluster of M1 circuit has a high probability occurrence with a large scale of biphasic dynamics but with a low scale of emergent dynamics (red square). While the sub-cluster of M2 circuit prefers to achieve a middle scale of biphasic dynamics but with a small or high scale of emergent dynamics (red and orange squares). M3 circuit sub-cluster is concentrated on attaining a small scale of biphasic dynamics, but with a high scale of emergent dynamics (red square). Therefore, despite containing the incoherent feedforward loop, the three minimal circuits exhibit divergence scales for achieving BE dynamics, providing potential diversity control strategies in regulating various biological functions.

Starting from the three minimal circuits, the 1,701 circuits that can achieve BE dynamics are generated by adding edges one by one. As a result, a comprehensive atlas describing the topological evolution of three-node circuits and their corresponding probabilities for achieving BE dynamics are entirely presented in Figure 6E. The topological complexity (the number of interaction terms/edges E) of the atlas increases from bottom to top, and the probability of a circuit of the same complexity to achieve BE dynamics decreases from left to right. The connectivity in the global atlas of topological evolution could supervise any one interaction to enhance or resist the circuit fulfillment function.

If an added edge improves the probability for achieving BE dynamics of the minimal circuit, such an edge is functionally significant^38, 46^. The statistical result of stochastically adding edges to improve the probability of achieving BE dynamics based on the minimum motif M1 is shown in Figure 6F. The result indicates that adding the self-activation of node C significantly increases the probability for inducing BE dynamics. The probability is decreased by adding inhibition, but is increased through considering the promotion of node C on node A. Based on the structure of adding node C self-activation and the promotion of node C on node A, the probability reaches the highest while further considering the inhibition of node B on node A and self-inhibition of node B in M1. Positive feedback loop is proved to play decisive roles for realizing multi-stable states in biological systems^39–41^. To discuss the structure of M1 for robustly achieving coexistent dynamics, six terms which could form positive feedback loops in motif M1, are individually (diagonal node) or integratively (non-diagonal node) added (Figure 6G). The proportion of the system with BEC dynamics based on the achieved BE dynamics system is calculated, showing that the contribution of node C self-activation is significant, and the proportion of system with coexistent dynamics is the highest when the promotion of node C on node-A is added.

Taken together, for robustly achieving BEC dynamics, the structure shown in Figure 6H should be the optimal circuit topology for M1, where red lines represent the essential edges while blue lines are the regulatory edges. Actually, the experimentally determined RIP1-RIP3-C8 circuit (Figure 1B) is highly consistent with the screened optimal circuit, revealing the precise design strategy of the biological system in controlling cell death. The optimal structure for circuits M2 and M3 are also discussed and the corresponding results are shown in Figures S7B and S7C.

**Figure 6.**
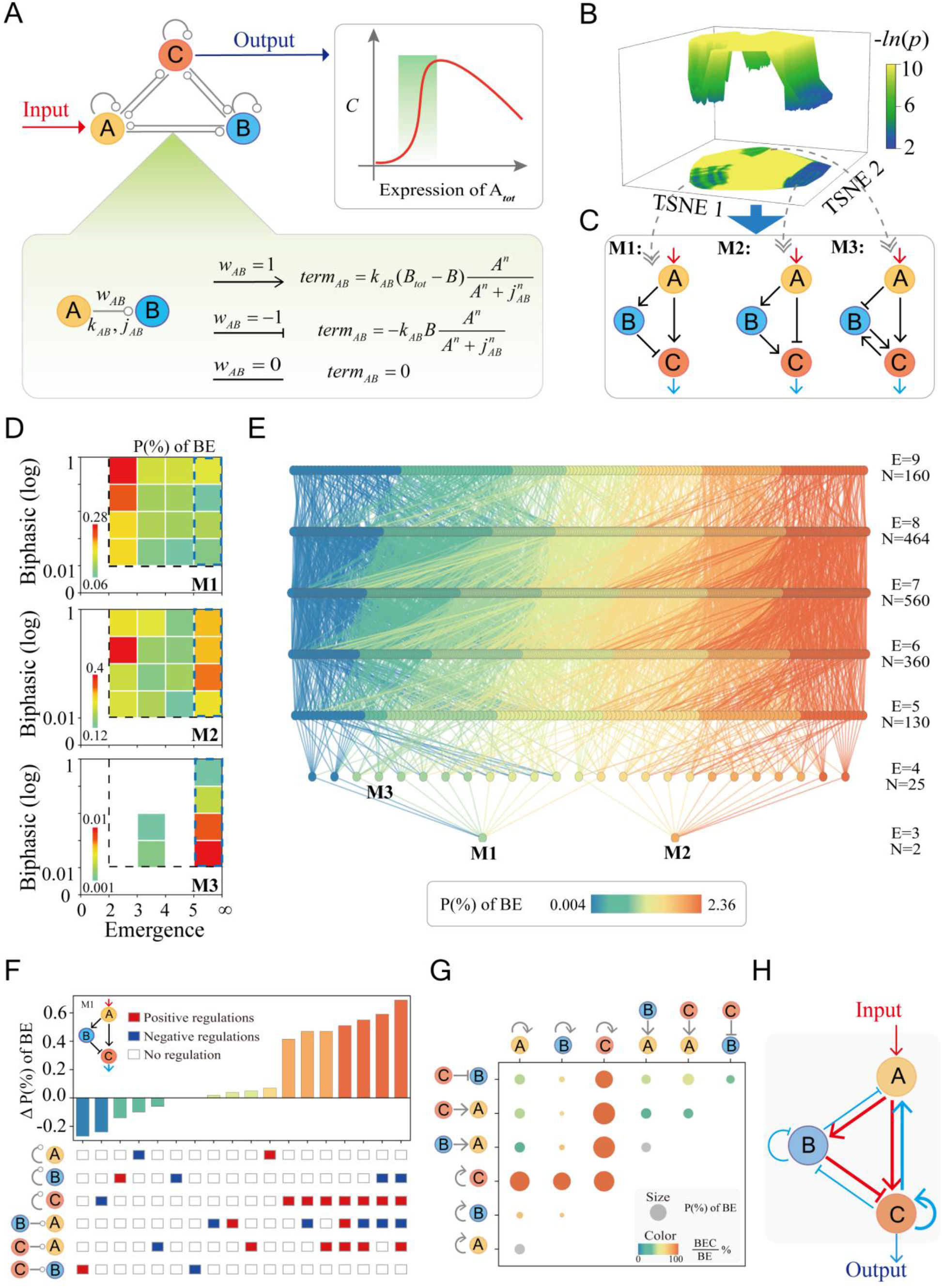
Topological exhaustive method reveals a complete atlas of achieving BE dynamics in three-node circuit. (A) Illustration of the structure screening procedure from mapping topology to function of asymmetric directed three-node circuit. A, B and C are the input node, regulatory node, and output node, respectively. There are three kinds of edges between any two nodes, *w* = 1 means promotion, *w* = -1 means inhibition, and *w* = 0 means no interaction. (B) Topological landscape of 4,698 coupling matrices in a 2D topological space. The well depth represents the probability of a sub-cluster achieving BE dynamics. (C) The minimal circuit of the sub-clusters that corresponds to the three wells in the topological landscape. (D) Probability distributions of the three minimum circuits are mapped into the biphasic dynamics and emergence 2D scale spaces. (E) A global atlas of 1,701 circuits that enable topological evolution of three-node circuits for achieving BE dynamics. (F) Probability statistics of BE dynamics that can be achieved by adding edges based on circuit M1. (G) The proportions of systems that with BEC dynamics when the six positive feedback edges are added to M1 individually (diagonal node) or in combination (non-diagonal node). (H) The determined optimal circuit for robustly achieving BEC dynamics.

## Discussion

Crosstalk between pathways is easy to understand, but why a given end-result is eventually reached is often a puzzle. As a key molecule in TNF signaling, RIP1 is required for inducing necroptosis^47^. While the inhibitory function of RIP1 is also demonstrated in mouse genetic studies^48, 49^. How RIP1 regulates dynamics of downstream substrates in determining specific cell death outcomes is a long-standing question^50, 51^. We previously quantified the RIP1-induced biphasic dynamics of necroptosis and further deciphered the control mechanisms^12^. However, the complexity of the system and large number of parameters involved in previous model limit the generalizability of conclusions and obscure the underlying mechanisms. To determine the topology and regulatory mechanism for necroptosis biphasic, emergent, and coexistent dynamics, we proposed a circuit cell death model of the TNF signaling based on previous studies^12^ and our experimental data (Figure 1). RIP1-RIP3-C8 incoherent feedforward loop is determined for achieving biphasic dynamics with emergence, while the positive feedback loop of RIP3 on RIP1 is required for death mode coexistence (Figure 2D). Instead of exploring the mechanisms with specific models, random parameter analysis of the TNF circuit is also performed, identifying that the incoherent feedforward loop embedded with RIP3 self-activation is another effective structure for achieving BEC dynamics (Figure 5G). We attempted to explore whether there exist general circuit design principles for natural systems to execute BE dynamics by using the topological landscape (Figure 6B), bottom-up, and topological evolution (Figure 6E) strategies. Both two- and three-node circuits are systematically analyzed and only three minimal three-node circuits are identified finally, confirming that the incoherent feedforward loop is the essential module for BE dynamics.

**Figure 7.**
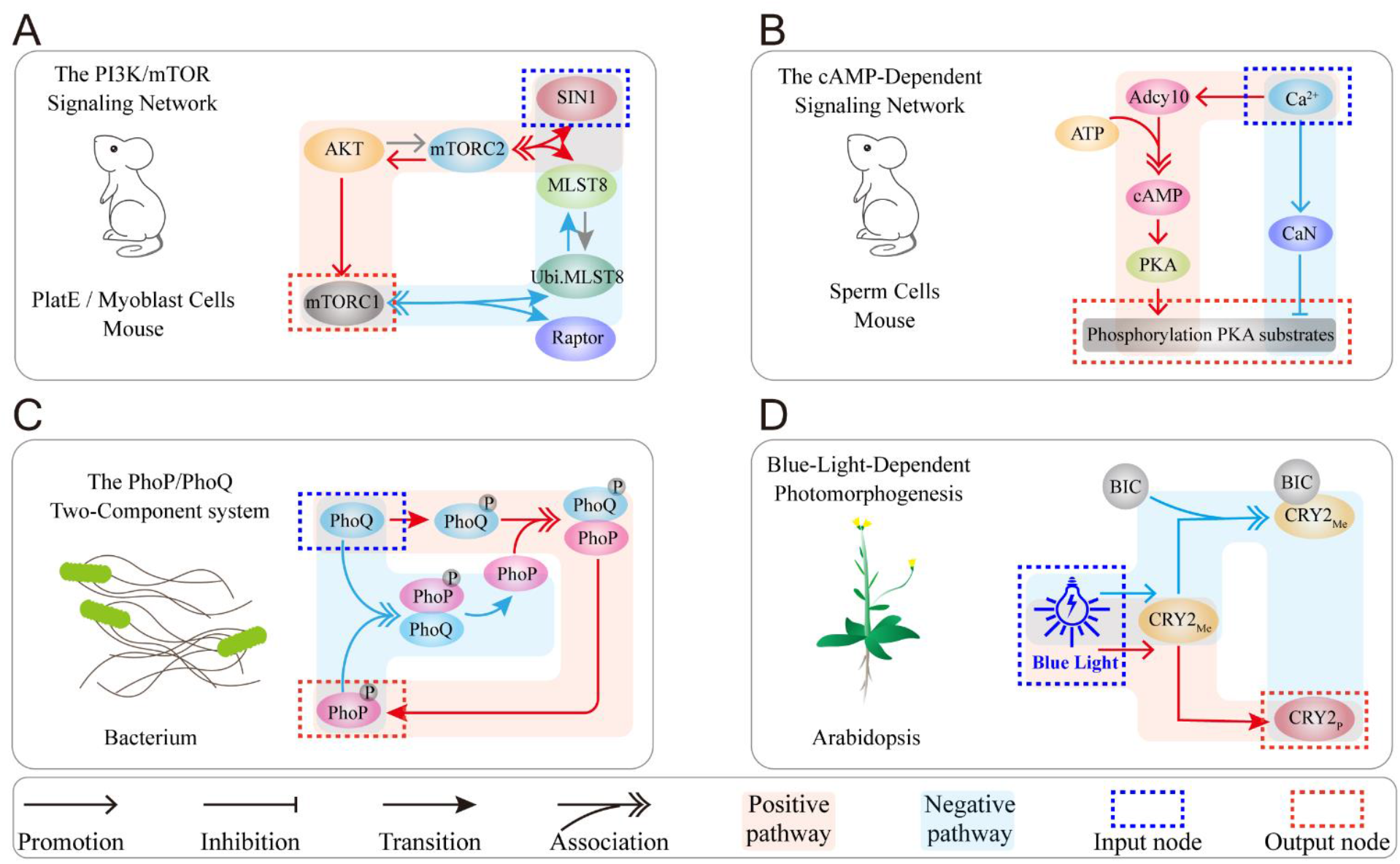
Examples of biphasic dynamics induced by incoherent feedforward loop in various cell types. Blue backgrounds represent the negative regulatory pathway, and red backgrounds in mTOR signaling. (B) Biphasic function of calcium ions in cAMP-dependent signaling. (C) The dual role of PhoQ in regulating PhoP phosphorylation in bacterial two-component system. (D) Biphasic response of cryptochrome photoreceptor 2 (CRY2) to photomorphogenesis in Arabidopsis.

Biphasic dynamics have been observed to drive essential biological processes in all forms of life, including mammalian cells, plant cells, and even bacterial cells. Previous study reported that SIN1, a key mTORC2 subunit, biphasically regulates mTORC1 activity in Myoblast cells^52^. The dynamics is determined by the incoherent feedforward loop shown in Figure 7A. Low-dose SIN1 promotes mTORC1 by synthesizing mTORC2, whereas high-dose SIN1 over-depletes MLST8 resulting in mTORC1 decreases. Biphasic dynamics is also observed in cAMP signaling which is triggered by the incoherent feedforward loop^53^ (Figure 7B). Calcium ions positively regulate Adcy10 to promote cAMP synthesis and PKA activation in sperm flagellum, and also inhibit PKA by activating CaN (calcineurin). Besides the mammalian cells, we previously found the biphasic dynamics of PhoP phosphorylation regulated by PhoQ in bacteria^7^. The incoherent feedforward loop exists in the PhoP/PhoQ signaling as well (Figure 7C). On the one hand, PhoQ promotes PhoP phosphorylation through binding to PhoP. On the other hand, excess unphosphorylated PhoQ also binds to phosphorylated PhoP to dephosphorylate PhoP. Our former study also observed the biphasic dynamics of CRY2 controlled by blue light in Arabidopsis^9^, and the incoherent feedback loop is also hidden in the CRY2 signaling (Figure 7D). With the increase of light intensity, blue light not only promotes the transition of CRY2_Me_ to CRY2_p_, but also promotes the combination of CRY2_Me_ and BIC to form a complex to reduce the level of CRY2_p_. Taken together, the topology of these signaling networks suggests that our determined incoherent feedforward loop should be a generalizable design principle for robustly executing biphasic dynamics in biological systems.

Despite the complexity and diversity of cell signaling networks, their core module and central topology should be highly conserved. Understanding the general design principles to achieve specific functions in diverse biological systems is significant. Forward searching all two- or three-node circuits are effective to find essential structures for achieving functions such as adaptation, noise-attenuation, robust oscillation, *etc.*^4, 16, 42^. The biological systems frequently exhibit muti-functions at the same time. Unlike the previous studies that mainly focused on one or two biological functions, we identified the topological structure that can achieve three general functions, *i.e.*, biphasic, emergent, and coexistent dynamics in this work. Among the identified circuits, auxiliary interaction on M1 motif that increase the probability of functional achievement are consistent with the experimentally observed RIP1-C8-RIP3 structure (Figures 2D and 6C), suggesting that the biological systems are naturally optimal structures. Of course, topological exhaustivity is also a powerful approach for predicting interaction in biological systems that have not been experimentally observed. In our analysis, the three identified circuits (M1, M2, and M3) seem to exhibit divergence scales for achieving BE dynamics (Figure 6D). While the intrinsic differences among these circuits are not captured. Further studies are still needed to systematically compare the general principles of these incoherent feedforward loops in exerting biological functions.

Cell states correspond to the attractors of the dynamical system, while potential landscape captures the dynamical principles of cell state transitions through providing a global characterization and stability measurement^54–56^. Potential landscape allows the targeted exploration of fundamental features and switching strategies of cell fate decision processes, and their application deepens our understanding of biological functions. Most recently, a new cell-aging fate induced by overexpression of the lysine deacetylase Sir2 was found by using this approach^57^. Besides, an unexpected observation of the lineage specifiers that can facilitate reprogramming and replace reprogramming factors of a corresponding lineage-specifying potential, was successfully clarified with landscape analysis as well^58^. Here, our study quantitatively provides the stochastic dynamics, the global nature, and the kinetic transitions of the cell death signaling. This is the first landscape discussion of necroptosis signaling to investigate the regulation of death mode switching. We systematically explored how the system structure changes the volume of the valleys, potentially helping to develop therapeutic strategies for death control. However, while employing the landscape theory, it is still difficult to use Fokker-Planck equation to solve the evolution probability of high-dimensional complex system. Although it has been proven effective to coarse-grain a high-dimension system into a low-dimension^59^, the curse of dimensionality exists objectively. Thus, deep learning method, truncated moment equations, partial self-consistent mean field approximation, and trajectory density should be developed and further considered in future study^59–62^.

For living systems with nonequilibrium multi-stable states, the essence of state switching is violating the principle of detailed balance that occurs at the cost of increasing entropy^63^. However, the complexity and only partial accessibility of living systems severely limit the inference of crucial thermodynamic quantities, like the entropy production. Previous studies mostly considered coarse-graining as a mapping method to simplify the complex systems to the reduced Markov networks. Recent studies also sought to measure the rate of entropy production by estimating the probabilities of forward and reverse trajectories in sufficiently long time series data. These theoretical explorations provided groundbreaking insights into understanding the central dogma, cells sense through receptors, and so on^37, 64–66^. In this study, Shannon entropy is introduced for the first time to measure the uncertainty of cell death mode transition. Information entropy presents a possible paradigm for understanding the transformation of energy and information in cells to perform fate decisions, and further consideration of the relationship between the information cost and the free energy cost of nonequilibrium systems is also urgently needed. These analyses will provide novel insights into the role of ‘Maxwell’s demon’ in fate decisions in living systems^67, 68^. However, due to the macroscopic limitation of complex living systems, our work assumes that the state transitions on the observed degrees of freedom are equivalent to the state transitions of all degrees of freedom of the system. We cannot determine whether there are other state transitions based on partial observations. The inference of information and energy associations in living systems is still an obvious and enormous challenge.

## Methods

### Cell line and cell culture

Mouse fibrosarcoma L929 were obtained from ATCC. RIP1 KO, RIP3 KO, L929, TRADD KO and Caspase-8 KO L929 cells were generated by TALEN or CRISPR/Cas9 methods. The knock-out cells were determined by sequencing of targeted loci and immunoblotting of the expression of respective proteins. All cells were maintained in Dulbecco’s modified Eagle’s medium (DMEM), supplemented with 10% fetal bovine serum, 2 mM L-glutamine, 100 IU penicillin, and 100 mg/ml streptomycin at 37°C in a humidified incubator containing 5% CO2. The target sites were designed as follows: RIP3: “CTAACATTCTGCTGGA”; RIP1: “AACCGCGCTGAGTGAGTTGG”; TRADD: “AAGATGGCAGCCGGTCAGAA”; Caspase-8: “GTGTTCAAATACATACGCCT”. All lentiviral-shRNAs were constructed into pLV-H1-EF1α-puro vector or pLV-H1TetO-GFP-Bsd following the manufacturer’s instruction (Biosettia). The indicated shRNA target sequences was: RIP1 shRNA: 5’-GCATTGTCCTTTGGGCAAT-3’.

### Reagents and antibodies

Mouse TNF-α were obtained from eBioscience (San Diego, CA, USA). Anti-RIP3 (dilution 1:1,000) and anti-MLKL (dilution 1:1,000) were raised using E. coli-expressed GST-RIP3 (287-387 amino acid), GST-MLKL (100-200 amino acids) and GST-FADD (full length), respectively. Anti-caspase-8 antibody (4790, dilution 1:1000) and anti-cleaved caspase-8 antibody (8592, dilution 1:500) were purchased from Cell Signaling Technology. Anti-p-RIP3 antibody (ab222320, dilution 1:500) and anti-p-MLKL antibody (ab196436, dilution 1:1,000) were purchased from Abcam. Anti-Gapdh antibody (60004-1-Ig, dilution 1:2,000) were from Proteintech. Anti-RIP1 antibody (610459, dilution 1:1,000) was from BD Biosciences.

### Immunoprecipitation and western blotting

Cells were seeded in a 100 mm dish, grew to reach confluency. After stimulating, cells were washed by PBS for three times and then lysed with lysis buffer (20 mM Tris-HCl, pH 7.5, 150 mM NaCl, 1 mM Na2EDTA, 1 mM EGTA, 1% Triton X-100, 2.5 mM sodium pyrophosphate, 1 mM β-glycerophosphate, 1 mM Na3VO4) on ice for 30 min. Cell lysates were then centrifuged at 20,000 g for 30 min. The supernatant was immunoprecipitated with anti-Flag M2 beads at 4°C overnight. After the immunoprecipitation, the beads were washed three times in lysis buffer and the immunoprecipitated proteins were subsequently eluted by SDS sample buffer with 0.15 μg/μL 3×Flag peptide.

### Microscopy imaging of cell death

To examine cell death morphology, cells were treated as indicated in 12-well plates or 35-mm glass bottom dishes for image capture. Static bright-field images of cells were captured using Zeiss LSM 780 at room temperature. The pictures were processed using Image J or the ZEN 2012 Image program.

### Noise and potential landscape

There is randomness in the procedure of biochemical reactions in cells, including intrinsic randomness, that is, molecular noise, and thermal fluctuations in the biochemical environment^69, 70^. For simplicity, we add a noise term, *σdξ*, to the OEDs of the deterministic model and assume that the noise intensity is correlated with the protein level. *ξ* represents for white Gaussian noise, and the statistical properties satisfy <*ξ_i_*(*t*)> = 0 and <*ξ_i_*(*t*) *ξ_i_*(*t’*)> = 2*σδ*(*t*-*t’*).

The global dynamics of a stochastic system with noise are given by the potential landscape. The stochastic dynamics of cell death fate decision system could be characterized by the generalized Langevin equation *dx_i_*(*t*)/*dt* = *F*(*x_i_*) + *ξ*(*t*), where *x* represents the concentrations of the proteins and *F* is the driving force. The Fokker-Planck equation describes the evolution of the probability density *p* in the state space, as following: ∂*p*(*x_i_*, *t*)/∂*t* = -Σ∂(*F*(*x_i_*) *p*(*x_i_*, *t*))/∂*x_i_* + *D_i_*Σ∂^2^*p*(*x_i_*, *t*)/∂*x_i_*^2^. Since the dimensionality of the model limits the direct access to the probability density through the evolution of the Fokker–Planck equation. We use the Bernoulli experiment numerical method to replace the steady-state probability distribution with the trajectory density distribution in the phase space. Specifically, we divide the two-dimensional phase space of RIP3 and C8 into 200×200 lattices and assign 10,000 sets of random initial conditions to the stochastic differential equations. After a long enough evolution, 10,000 trajectories can be obtained in the phase space, and the number of trajectories in each lattice is counted and the density is calculated^58^.

### Identification of biphasic dynamics with emergence

In this study, the system depends on normalized TNF stimulation to be activated, the strength of which is a random value in the range (0,1]. The levels of C8 and pRIP3 are also dependent on the upstream signaling molecule RIP1, and the knockdown of RIP1 means that the signal could not be transmitted to downstream signaling molecules. In numerical simulations, the scale H of the biphasic kinetics is dependent on the peak pRIP3 level and the level when RIP1 is not knocked down (deterministic model). The expression of RIP1 is fixed as a control parameter, discretized with a step size of 0.02 in the normalized parameter space. First, a two-dimensional array of RIP1-dependent pRIP3 level and an index of RIP1 expression corresponding to pRIP3 peak are obtained through time evolution. Second, the elements in the two-dimensional array must satisfy the characteristics of monotonically increasing on the left of indexpRIP3peak and decreasing monotonically on the right. Finally, the pRIP3 level at 100% RIP1 expression must be 0.01 lower than the *pRIP3_peak_*. The emergent behavior should satisfy the level of pRIP3 increased by not less than 50% when RIP1 expression continuously increases by 10%.

### Explanation of topological exhaustive method

Here we seek to uncover the minimal core motifs that enable biological networks to achieve biphasic and emergent kinetics. The study of functional motifs in complex biological networks based on node directed networks has been widely reported^4, 38, 42^, and thus is also applicable to the achievement of biphasic and emergent kinetics for more complex biological networks embedded with the minimal core motifs we found.

For the two-node directed network, each topology corresponds to a 2×2 coupling matrix, and each coupling edge could be assigned to 1 (promoting), -1 (inhibiting), and 0 (no interaction). There are theoretically 3^4^ (=81) topologies. However, the heterogeneity of input and output nodes was considered in this study, the activation of the output node depends on the input node, and the control parameter is fixed to the protein (gene) expression amount represented by the input node. Therefore, the coupling edge of the input node to the output node can only be fixed to 1, and finally only 27 topologies are considered. For a three-node directed network, a control node is introduced, and the system also relies on the activation of input nodes. The constraints of interaction include 1) the output node must have a promoting action from the input node or regulatory node; 2) the regulatory node must have a promoting action from the input node or output node; 3) one of the input nodes and output nodes must be promoted or inhibited by the regulated node. A class of motifs is eliminated from all motifs that meet the above three conditions, and their output node and regulatory node promote each other but are not promoted by the input node. The theoretical 3^9^ (=16983) topologies are reduced to 4,698.

### Parameter ranges selection and sampling

In our study, the parameter ranges of the computational models were consistent with those in previous publications of similar studies^4, 38, 71^. For each topology, 50,000 parameter sets are uniformly sampled using Latin hypercube sampling, with parameters ranging from *k*∼0.1-10 (logarithmic scale), *j*∼0.001-100(logarithmic scale), *d*∼0.01-1(logarithmic scale), n∼1-4(integer scale), stimulation signal∼0-1.

## Data availability

The data of this work are available from the corresponding author upon reasonable request.

## Code availability

The key codes for this work are deposited to GitHub at https://github.com/XMU-Xu/ANscn.

Other reasonable requirements can be obtained from the corresponding author.

## Acknowledgements

We greatly thank PhD. Ming Yi (School of Mathematics and Physics, China University of Geosciences, Wuhan) and PhD. Peng Ji (Institute of Science and Technology for Brain-Inspired Intelligence, Fudan University, Shanghai) for helpful discussion of this paper. This work was supported by the National Natural Science Foundation of China (Grants Nos. 12090052 and 11874310), the Ministry of Science and Technology of the People’s Republic of China under grant nos. 2021ZD0201900 and 2021ZD0201904, and the Fujian Province Foundation (Grant No. 2020Y4001).

## Author contributions

Fei Xu conceived the idea, developed the algorithms, analyzed the data and wrote the paper. Xiang Li conceived the idea, analyzed the data and wrote the paper. Rui Wu performed the experimental analysis. Hong Qi, Jun Jin, Zhilong Liu, Yuning Wu, Chuanshen Shen and Hai Lin helped to analyze data. Jianwei Shuai revised the paper and supervised the project. Fei Xu and Xiang Li contributed equally to this work.

## Supplemental Information

### Computational modeling

Ordinary differential equations are built to describe the evolutionary dynamics of cell death signaling mediated by TNF^71, 72^. For example, the interaction of RIP3 phosphorylation by pRIP1 is:

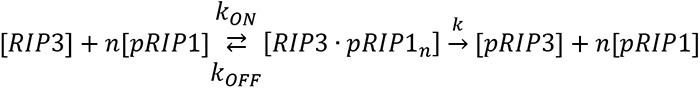

The equation of the intermediate complex [pRIP1·RIP3] is described as follows:

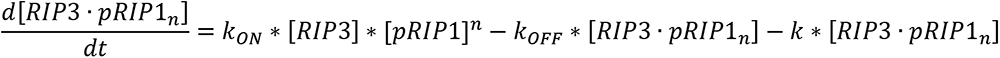

At steady state

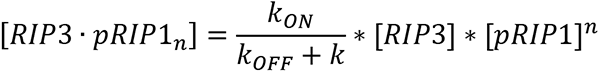

Assuming the binding between proteins is independent, and the dissociation rate of the complex is much larger than the binding rate.

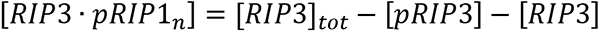

With the normalized total amount of proteins, the rate of pRIP1-mediated phosphorylation of pRIP3 is described as follows:

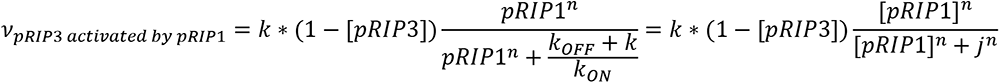

The complete equations of deterministic TNF circuit model are presented below:

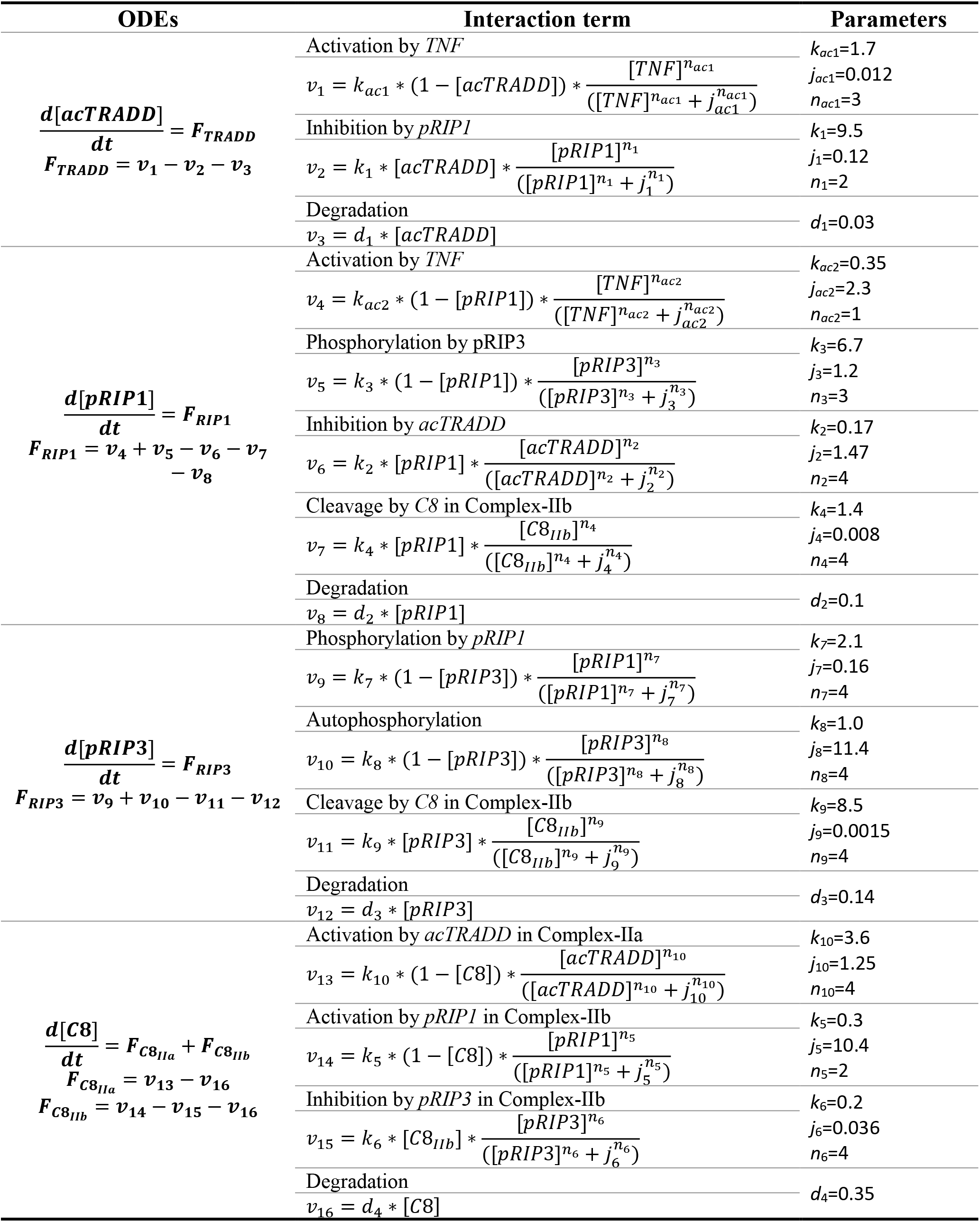

### Kinetic parameters estimation

Table of public experimental data sources for parameter estimation^12^.

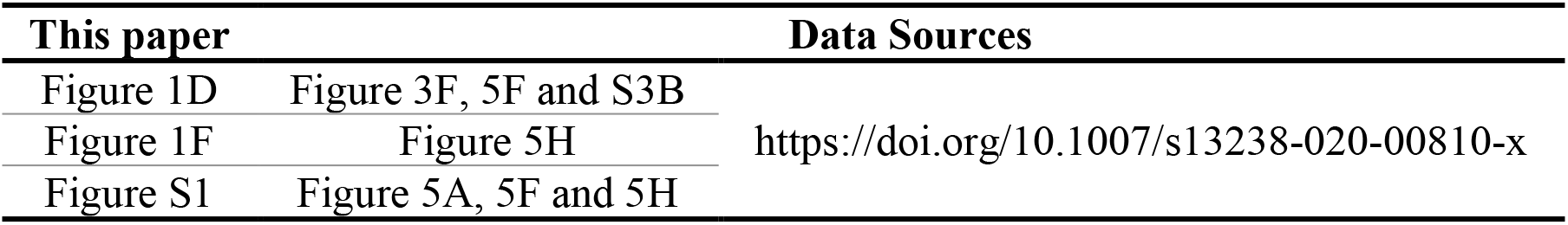

### Supplemental Figures

**Figure S1.**
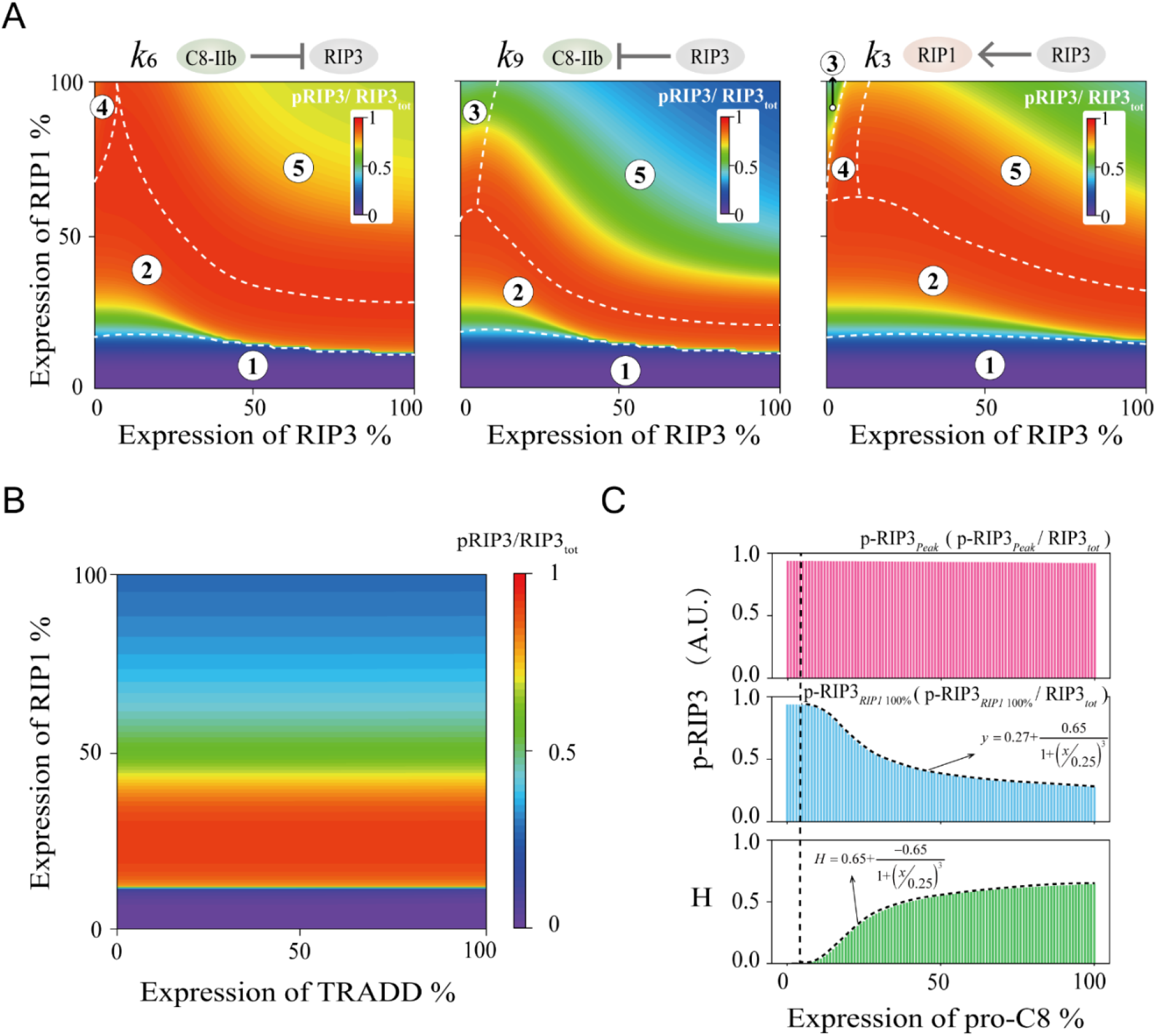
(A) The relative level of pRIP3 in the RIP3-RIP1 phase space when the terms of k6 (inhibition of C8 on pRIP3), k9 (the inhibition of pRIP3 on C8), and k3 (the positive feedback of pRIP3 on RIP1) are reduced 10-fold, respectively. (B) The relative level of pRIP3 in the TRADD-RIP1 phase space. (C) The variation of pRIP3Peak, pRIP3RIP1_100%, and H with pro-C8 expression level increases.

**Figure S2.**
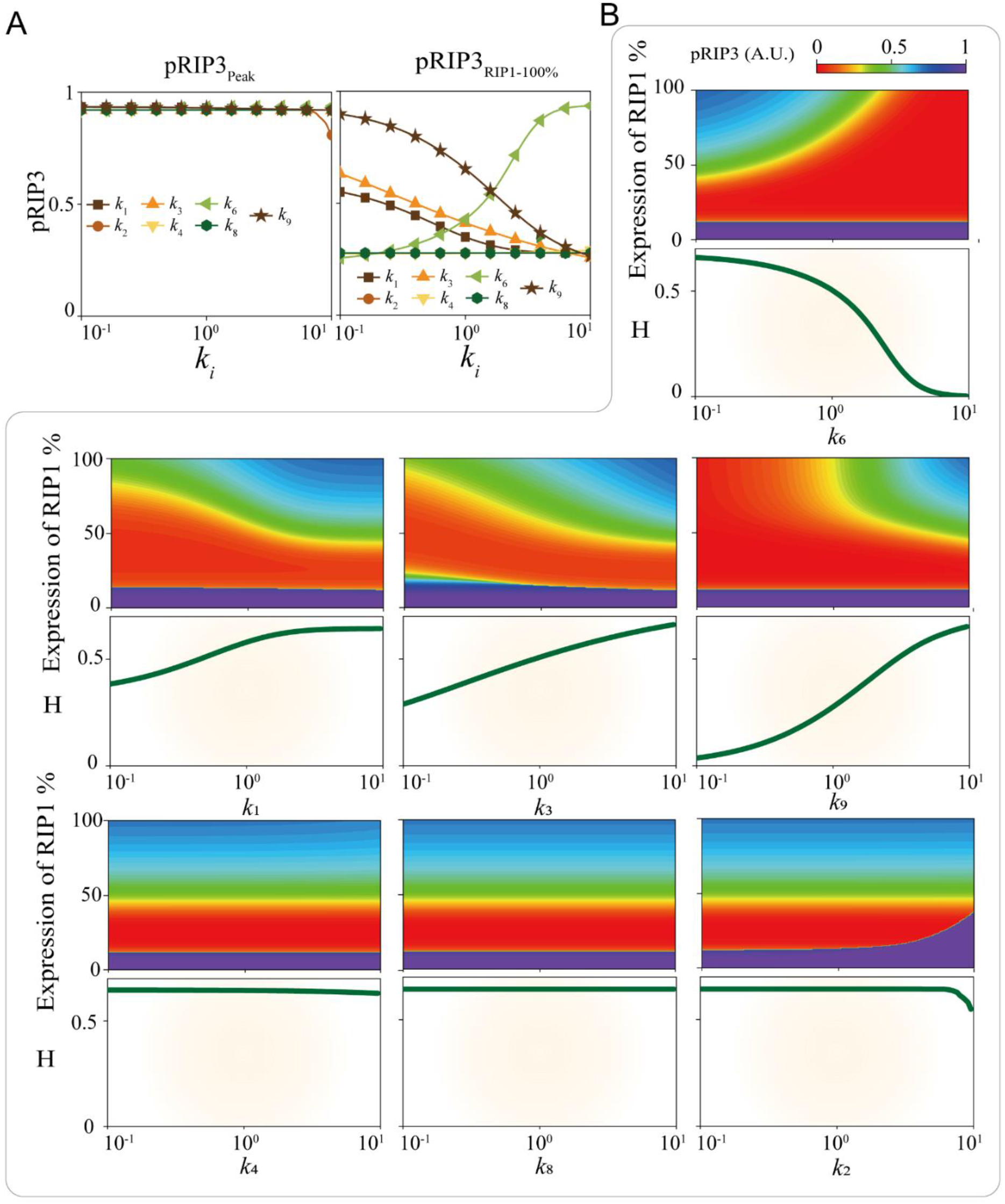
(A) Parameter sensitivity of other seven terms in modulating *pRIP3_peak_* and *pRIP3_RIP1_100%_*. (B) Analysis of the seven terms regulations on pRIP3 and the scale of biphasic dynamics H.

**Figure S3.**
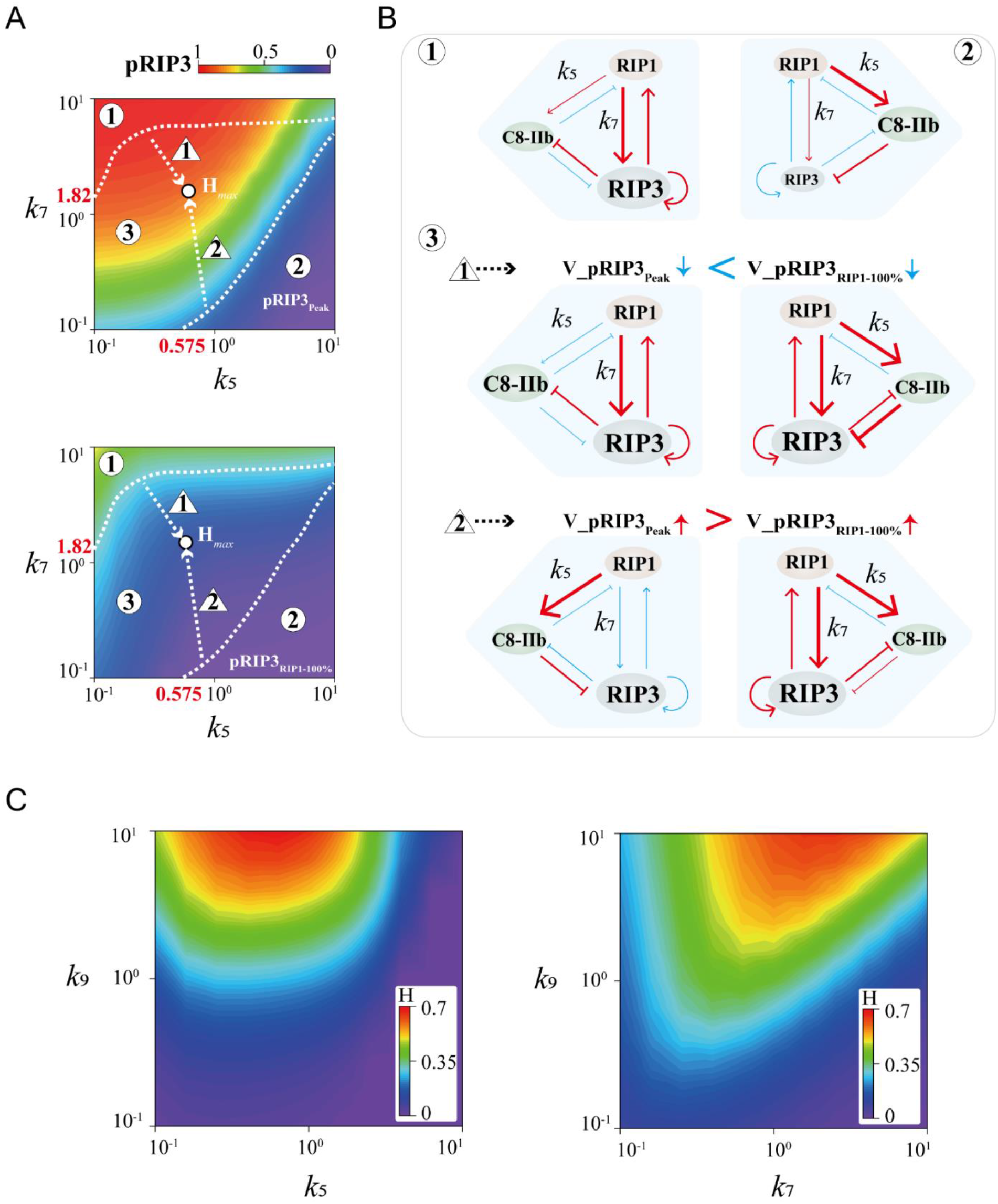
(A) Levels of *pRIP3peak* and *pRIP3RIP1_100%* in the *k*5-*k*7 parameter space, and the phase plane is decomposed into three regions and two processes. (B) Mechanistic analysis of *k*5 and *k*7 Bell-shaped regulation on pRIP3 biphasic dynamics. In regions 1 and 2, two terms, activation of RIP3 by RIP1 and activation of C8 by RIP1, play the dominant role, respectively. Their corresponding *pRIP3peak* and *pRIP3RIP1_100%* are both high or both low, resulting in small scales of biphasic dynamics. The decline rate of *pRIP3peak* in process 1 is lower than that of *pRIP3RIP1_100%*, and the increase rate of *pRIP3Peak* in process 2 is greater than that of *pRIP3RIP1_100%*. (C) Phase diagram of H in *k*5-*k*7 and *k*7-*k*9 parameter spaces.

**Figure S4.**
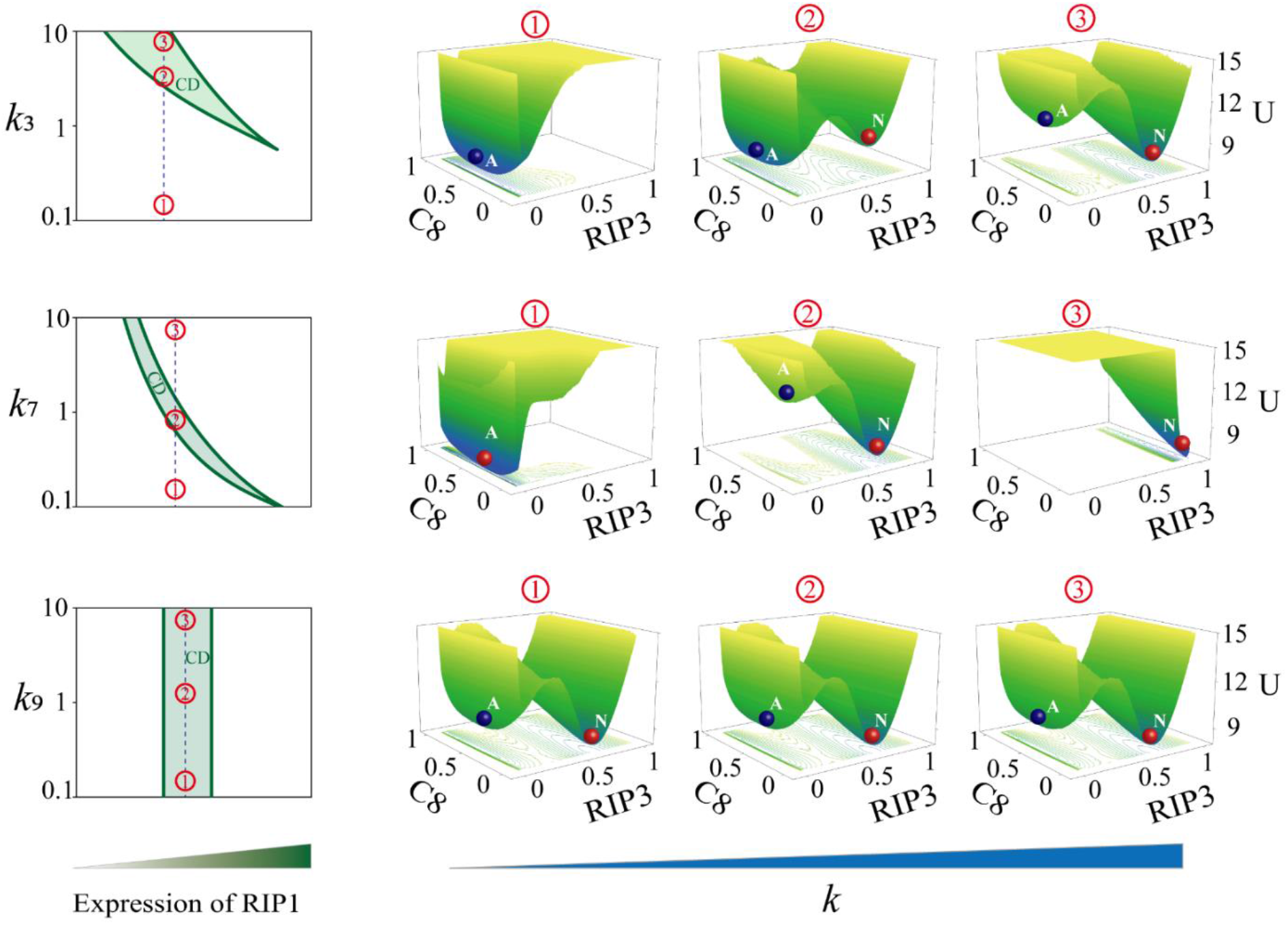
Phase diagram of the system stability on the co-variation of interaction term (k3, k7, and k9) and the expression of RIP1. The green shaded region indicates the coexistence of apoptosis and necroptosis. The potential energy landscape in C8-RIP3 phase space of the cell death system at three typical values fixed for each parameter.

**Figure S5.**
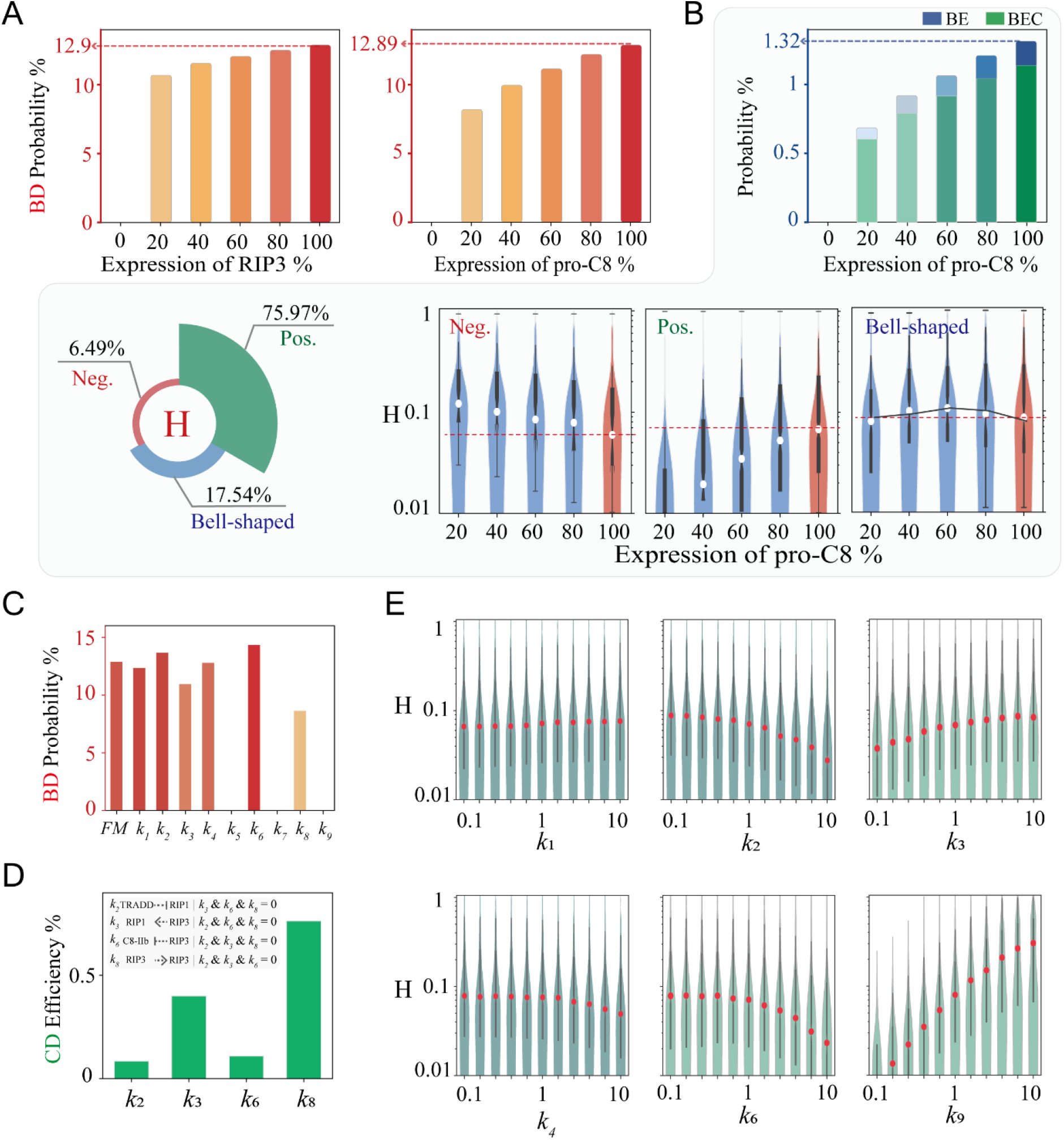
(A) Random circuit analysis with five representative RIP3 and pro-C8 expression levels to count the probabilities for achieving pRIP3 biphasic dynamics. (B) Random circuit analysis with five representative pro-C8 expression levels to count the probabilities for achieving pRIP3 BE and BEC dynamics, and the statistics of the regulatory behavior of RIP3 on the scale of biphasic dynamics H. (C) The probability of the system achieving biphasic dynamics when all interactions are blocked, respectively. (D) Contribution of all the positive feedback loops in circuit to achieve coexistence dynamics. (E) Statistics of the regulation of other six terms on H.

**Figure S6.**
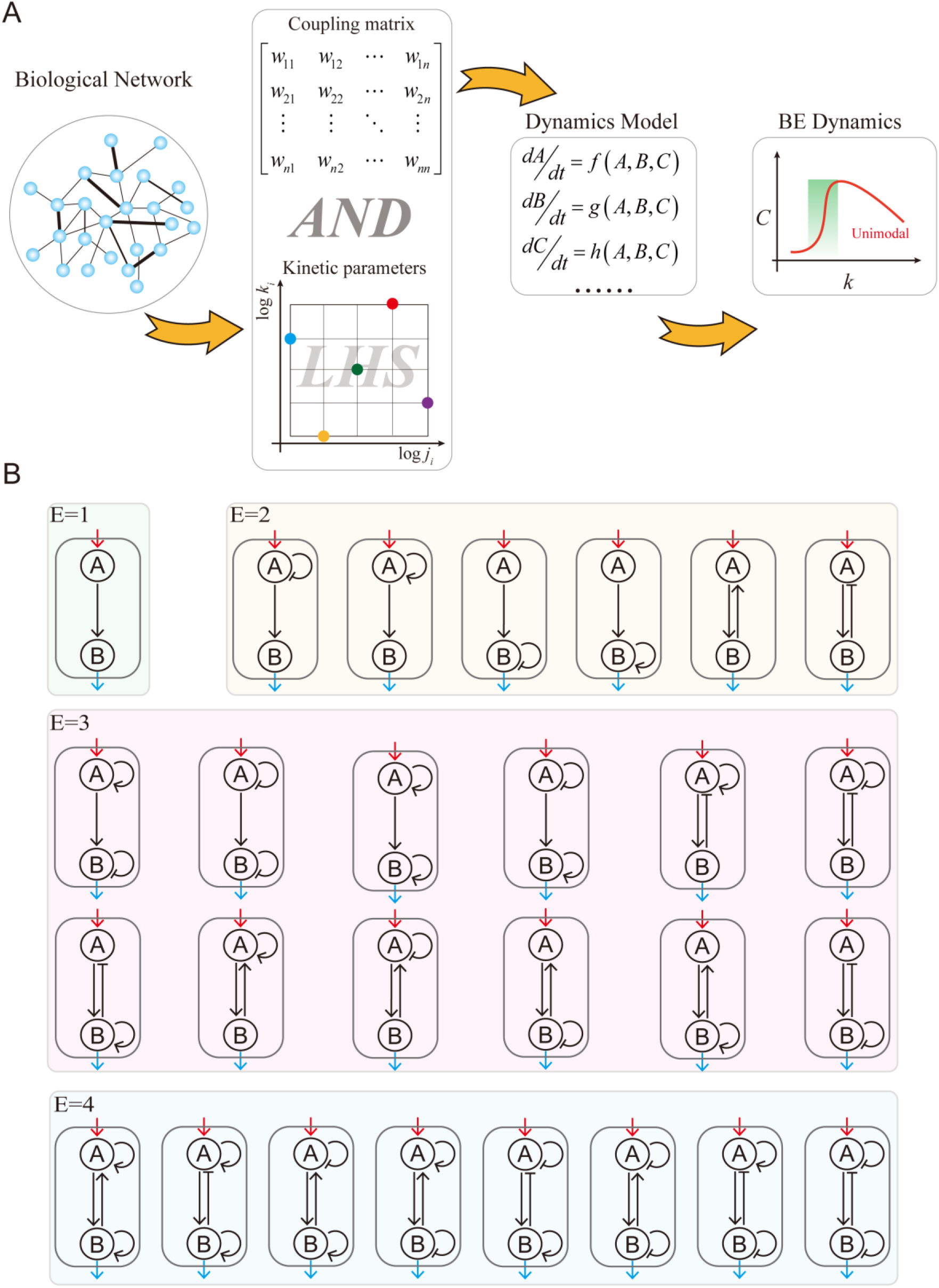
(A) Workflow for topology-to-function mapping of BE dynamics. (B) The 27 two-node motifs are classified according to the number of terms (connecting edges E).

**Figure S7.**
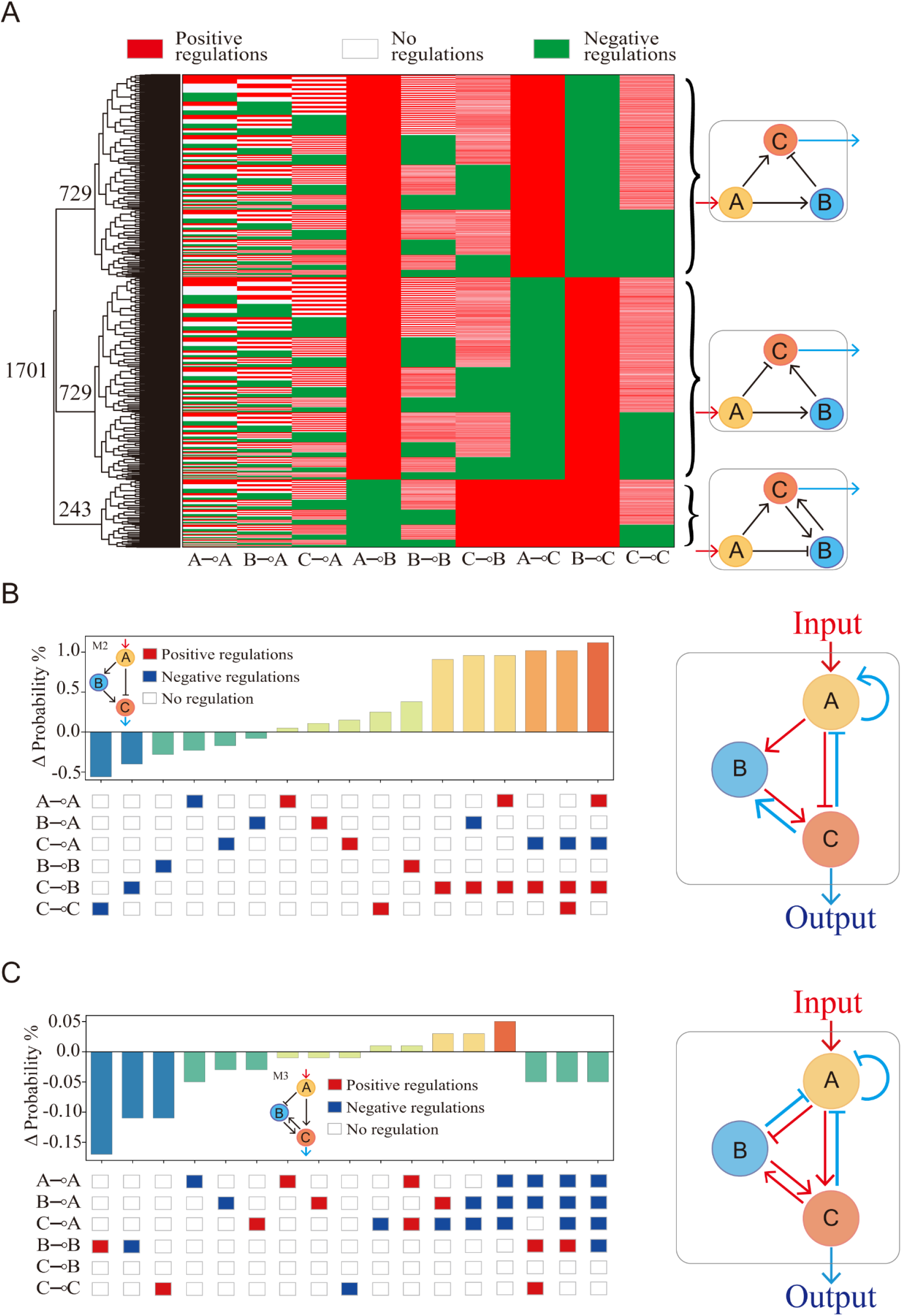
(A) Clustering of the 1,701 three-node circuits that can achieve BE dynamics. The core circuits associated with each of the sub-cluster are shown on the right. (B) and (C) Probability statistics of BE dynamics that can be achieved by randomly adding edges based on circuit M2 and M3.

